# Long-read genome sequencing reveals complex variability in lentiviral provirus insertions in deeply characterized Clonal CD19 CAR-T vector copy number reference cell lines

**DOI:** 10.64898/2026.06.15.731627

**Authors:** Zhiyong He, Jennifer H. McDaniel, Linhua Tian, Mahir Mohiuddin, Ningchun Xu, Lili Wang, Justin M. Zook, Hua-Jun He

**Affiliations:** Materials Measurement Laboratory, National Institute of Standards and Technology, 100 Bureau Dr. Gaithersburg, MD 20899, USA

**Keywords:** Chimeric antigen receptor (CAR)-T cell, lentiviral vector, vector copy number, integration site, reference material, long-read genome sequencing

## Abstract

Chimeric antigen receptor (CAR)-T cell therapy is an important therapy involving provirus insertions in the genome. Characterizing these insertions is important for understanding the safety and efficacy of cell products, but the sequence of these insertions is not fully characterized. We generate clonal CD19 CAR-T cell lines with one to five copies of the lentiviral provirus insertions. Vector copy number (VCN) was determined by droplet digital PCR (ddPCR), which revealed that most of the elements (LTR, Psi, RRE, CD19, and WPRE) were 1 to 5 or 6 copies per cell. DdPCR data also revealed that there was an additional copy of eGFP gene in VCN4 and VCN5 cell lines. To fully characterize the sequences and locations of these insertions, we use short- and long-read whole genome sequencing as well as digital PCR and flow cytometry. Long-reads enable full resolution of each insertion, and we find that of 10 insertion events, 3 have the expected insertion sequence, 2 differ from the expected only in small variants, 3 have structural abnormalities, and 2 are small partial insertions missed by most other approaches. One particularly important structural abnormality resolved only by long-reads is a 724 bp deletion of the EF1α promoter disrupting expression of the CD19 CAR. Standard short-read and ddPCR approaches miss this deletion due to this commonly used promoter being in the unengineered human genome. These results demonstrated that these cell lines are suitable VCN reference standards for 1 to 5 or 6 copies and highlight the utility of long-read sequencing in characterizing both quantity and quality of insertions in lentiviral-engineered cells.

## Introduction

Chimeric antigen receptor (CAR)-T cell therapy is a revolutionary novel cancer therapy and has shown promises in both leukemia and solid tumor ^1-5^. A significant milestone for CAR-T cell therapy was FDA approval of Tisagenlecleucel and Axicabtagene Ciloleucel, two CD19-specific CAR-T cell products^4,5^. Currently, there are 7 CAR-T cell therapy products approved by the US Food and Drug Administration (FDA) for treating blood cancers such as acute lymphoblastic leukemia (ALL), large B-cell lymphoma, and multiple myeloma (https://www.fda.gov/vaccines-blood-biologics/cellular-gene-therapy-products/approved-cellular-and-gene-therapy-products). Many other cancer related cell surface markers were targeted by CAR-T cell therapy in preclinical and clinical stages, such as CD22^6^, HER2^7^, and EGFR^8^, inspired by the success of these CAR-T cell therapies. A CAR molecule typically contains four functional domains: an extracellular antigen-binding domain, a spacer, a transmembrane domain, and an intracellular activation domain^9^. The T cells for therapy (either autologous or allogeneic) are genetically engineered to express the CAR protein.

There are at least two main methods for engineering the CAR-T cells currently. One is using CRISPR/Cas9 to engineer the T cells at specific sites, but none are yet FDA-approved^10-13^. The commonly used method for CAR-T production of therapies approved by FDA is the lentivirus vector (LVV) transduction. LVV is a retrovirus vector derived from HIV, and able to insert the target gene into the cell genome^14^. Thus far, 141 CAR-T and 6 CAR-NK cell therapies utilize lentivirus as the gene delivery method (www.clinicaltrials.gov as of May 2026). Advantages of the LVV method include relatively reliable and efficient gene transfer. However, LVV insertion into the target cell genome is uncontrolled, which could lead to aberrant RNA splicing ^15^. High copies of LVV integration may also cause insertional mutagenesis and/or oncogenicity^16,17^. Multiple insertions in the same chromosome close to each other may cause the fragment between the insertions inverted or deleted through homologous recombination. Furthermore, the biological process of lentiviral integration (including reverse transcription of LV RNA genome and double-stranded proviral DNA insertion) is inherently error-prone ^14,18,19^. Therefore, to reduce safety concerns, the FDA recommended the inserted vector copy number (VCN) shall be less than 5 copies per genome. Further studies of a gene therapy for hematologic cancer identified^20^ insertions within oncogenes in patients with subsequent cancer^20^, which suggested that identification of LVV insertion sites is important for safety. Several case reports prompted an FDA investigation and longitudinal study, which did not find evidence directly linking CAR-T cells to these cancers, but is still early ^21,22, 23,24^.

Common measurement methods for integrated LVV copy number include qPCR, ddPCR and next generation sequencing (NGS), each with its own limitations. For example, an earlier report showed that the integrated VCN was often underestimated by qPCR method^25^. In addition, the assays developed based on each method vary from laboratory to laboratory. Targeted NGS of integration sites has been used to measure integration with short-reads^26^ and long-reads^27^, but these targeted methods did not measure the full insertion sequences. Long-reads have detected large structural variants at CRISPR-Cas9 engineered sites,^28^ leading the FDA to mention the use of long-reads or alternative methods for large variants (https://www.fda.gov/media/191966/download). Targeted long-reads identified HIV-associated internal proviral deletions, and a modified approach recently found promoter deletions in LVVs as well. ^19,29^ For these reasons, the community has called for reference samples for assay validation, optimization, and benchmarking^30,31^.

We previously developed NIST RGTM 10268 using a LVV containing only a cassette for GFP expression^32^. The clonal cell lines contained 0-4 copies of proviral DNA per diploid genome, and all insertion sites were identified^32^. Here, we developed an additional set of VCN candidate reference material for the following reasons. First, the actual lentiviruses used in the clinical setting such as cell and gene therapy usually have longer RNA genomes. Second, since higher VCN samples are more difficult to reach consensus measurement results ^33,34^, higher VCN clonal cell lines need be included in the candidate reference materials for benchmarking the measurements. In a World Health Organization (WHO) interlaboratory study, measurements of cells with an estimated VCN of 10.67 varied widely from 6.15 to 13.6, and few insertions sites were identified ^34^. This broad range highlighted the challenges of measuring VCN for higher copies of high-copy-number integrated vectors. Moreover, the FDA guidance recommends maintaining a VCN of fewer than 5 copies per cell. Third, there is no CAR expression cassette in the previous candidate reference materials for potential functional CAR cytotoxicity assays. Finally, the proviral DNA sequence of previous candidate reference materials could not be disclosed due to intellectual property restrictions.

To address this unmet need, we developed a set of clonal cell lines with 1 to 5 copies of integrated CD19 CAR LVV, and deeply characterized the integration sites using orthogonal methods. Precise sequences of each LVV insertion were measured with long-read whole genome sequencing, revealing functionally important structural anomalies in the LVV insertion sequences in the host genome relative to the expected provirus DNA sequence. These clonal cell lines could serve as reference materials for CAR-T cell VCN measurements, insertion sites identification, and identification of potential random insertions.

## Materials and Methods

### Establishment of CD19 CAR-T VCN standard cell lines

Anti-CD19 CAR-T lentivirus transfer vector was designed at NIST based on the 3^rd^ generation of lentivirus packaging system ^1^. CD19 CAR vector design was illustrated in Fig. 1A. Briefly, CD19 CAR coding sequence was driven by an EF1α promoter. A Wood chuck Hepatitis Virus Postranscriptional Regulatory Element (WPRE) sequence was placed downstream of CD19 CAR coding sequence. EGFP coding sequencing and Bleomycin resistant sequence were separated with a P2A sequence under the same PGK promoter. The transfer vector sequence was validated by Sanger sequencing. The lentivirus particles were packaged by VectorBuilder (Malvern, USA). Jurkat cell line (E6-1), purchased from ATCC (Cat# TIB-152, Manassas, VA), was transduced at different multiplicity of infection (MOI=1, 5, and 10) to generate the VCN cell lines. The transduced cells were then selected with a Bleomycin mass concertration of 10 μg/mL in RPMI-1640 growth medium supplemented with 10 % fetal bovine serum (FBS Thermo Fisher, Cat# A5670402) for selecting the positive integration for the first 3 passages and then with 5 μg/mL of Bleomycin. The selected positive single cells were deposited into each well of 96-well plates by using Cytena Single-Cell Printer (Cytena) to generate the clonal cell lines. The clonal cell lines were screened by droplet digital PCR (ddPCR) using RRE ddPCR assay ^35^ and S2 or RPL32 as reference gene. The VCN cell lines were then further expanded and characterized.

**Fig. 1.**
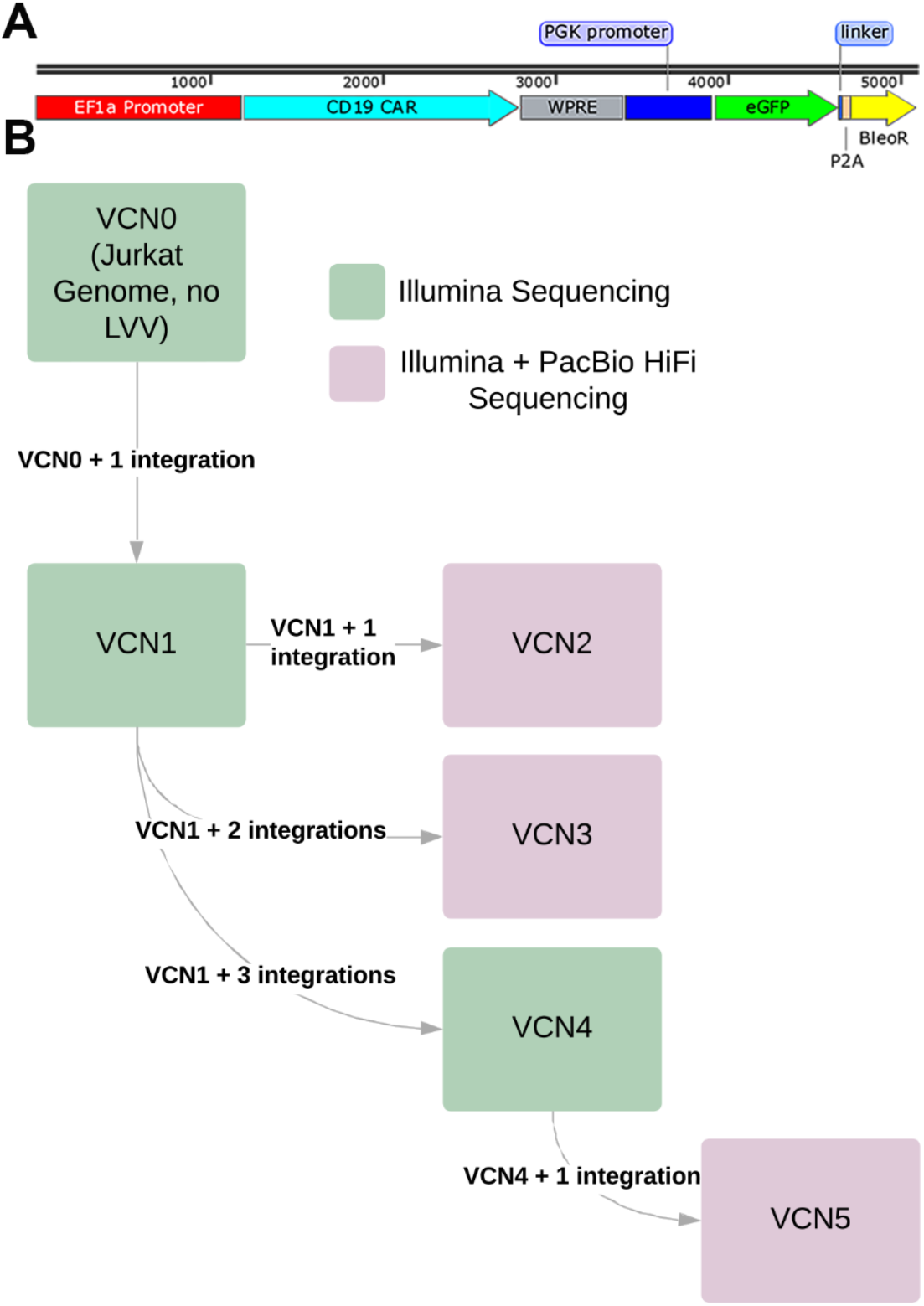
CD19 CAR vector design, and schematic view of VCN clonal cell lines generation strategy. A, Schematic view of CD19 CAR transfer vector design. B, Schematic view of VCN clonal cell lines generation strategy.

### Droplet digital PCR (ddPCR)

Integrated vector copy number (VCN) was measured by ddPCR using the common 3^rd^ generation lentiviral vector elements, LTR, Psi, RRE, and WPRE. The primer and probe sequences are listed in our previous publication ^35^. Additional target gene eGFP and CD19 CAR were also used to measure the VCNs. Bio-Rad ddPCR Supermix for probes (no dUTP) was purchased from Bio-Rad (Cat# 186-3024, Hercules, CA). Genomic DNA (gDNA) was extracted from each cell line using Quick-DNA Miniprep Kit (Zymo Research, Cat # D3024). Five to 20 ng of gDNA was used per ddPCR reaction. VCN was calculated as the ratio of lentiviral vector elements to reference gene (S2 and RPL32) determined with the Bio-Rad QX200 ddPCR system following manufacturer’s instructions.

### Flow cytometry analysis

The expression of eGFP and CD19 CAR in the clonal VCN cell lines was analyzed by flow cytometry. The cells (5 ×10^5^ cells per sample) were washed with PBS containing 10 % FBS (FACS buffer) once and resuspended in 100 μL of FACS buffer. The cells were then incubated with 2 ng of antibody, biotinylated anti-FMC63 scFV antibody (Acrobiosystems, Cat# FM3-BY45) on ice for 30 min. After 30 min, the cells were washed with 1 mL of FACS buffer once and resuspended in 100 μL of FACS buffer and 100 ng of APC conjugated streptavidin was added to each sample and further incubated on ice for 30 min. The cells were washed once with FACS buffer and resuspended in 300 μL of FACS buffer and then subjected to CytoFLEX LX Flow Cytometer. The experiment was repeated three times to confirm the repeatability of the results.

### Next generation sequencing (NGS)

The lentiviral vector integration sites in parental Jurkat cell line genome were determined by whole genome sequencing analysis. Genomic DNA extracted from each cell line was sent to Novogene for short-read Illumina library preparation and whole genome sequencing. Whole genome long-read PacBio sequencing was performed by Maryland Genomics. The raw sequencing data was analyzed at NIST to identify copy numbers of the integrated provirus sequence and their precise integration sites. Whole genome sequencing data was deposited at the NIH/NCBI/SRA under BioProject [PRJNA1282768](https://www.ncbi.nlm.nih.gov/bioproject/1282768) (Biosamples: SAMN49572894: VCN0, SAMN49572895: VCN1, SAMN49572896: VCN2, SAMN49572897: VCN3, SAMN49572898: VCN4, and SAMN49572899: VCN5.)

### NGS read alignment to reference genome

We used short and long-read whole genome sequencing (Illumina 2 × 150 bp and PacBio HiFi, N50 range of 18 751 to 22 133 bp) to identify LVV integration sites (IS) in the Jurkat cell genome. Reads were mapped to a modified reference genome (GRCh38-GIABv3_plusInsert-with-3primeLTR_flanks.fasta). Specifically, the expected 7374 bp LVV sequence was added as an additional chromosome to the GIABv3 (https://ftp-trace.ncbi.nlm.nih.gov/ReferenceSamples/giab/release/references/GRCh38/) GRCh38 human reference genome, which masks false duplications and adds decoy sequences from CHM13 ^36^.

Unlike the original LVV RNA sequence (7321 bases), the LVV sequence in our reference has identical 5’ and 3’ LTRs because during reverse transcription the U3 region of the viral genome is copied to the 5′ LTR (7374 bp) ^37^. Both long and short-reads were mapped to the modified reference using BWA-MEM (v0.7.17, https://arxiv.org/abs/1303.3997) with default parameters. The resulting alignment files (.sam) were then converted to .bam files, sorted and indexed with Samtools (v1.9) ^38^ view, sort and index, respectively. Duplicates were then marked with Picard (v2.18.29) MarkDuplicates (https://broadinstitute.github.io/picard/) for short reads. Alignment post processing steps used tool default parameters. Additionally, long-reads were mapped to the modified reference using pbmm2 (v1.13.1), a minimap2 ^39^ wrapper for PacBio reads, using default parameters. BWA alignments of Illumina and PacBio reads resulted in approximately 27 × coverage. Pbmm2 alignments of PacBio reads resulted in approximately 38 × coverage.

### Variant Calling

Variant calling was performed to confirm the expected number of integration sites (IS) in the Jurkat cell genome, identify the integration locations in GRCh38, and compare the actual inserted sequence to the expected LVV sequence. Insertions were identified using two distinct structural variant (SV) callers: Delly (v1.2.6) ^40^ for short-reads data and PacBio pbsv (v2.9.0, https://github.com/PacificBiosciences/pbsv) for long-read sequencing data. Variant calling was performed with the BWA short-read alignments and pbmm2 long-read alignments against the modified reference.

### Identification and confirmation of putative integration sites using NGS

To identify putative integration sites, Delly was used to identify translocations and breakpoints when variant calling with BWA aligned short reads resulted in passing SVTYPE= BND (breakend) on the LVV chromosome with ALT sequence identified on a different chromosome. Putative integration sites were then confirmed by manually curating sites using all alignments and variant calls with the Integrated Genome Viewer (IGV v2.16.2). For a given VCN, the VCN alignments, variant calls and modified reference genome were loaded into an IGV session. VCN0 alignments were also loaded in to serve as a negative control, as no integrations are expected in this sample. All alignments to the LVV chromosome were first examined to identify reads that were partially or fully aligned to the expected LVV sequence. BWA short-read alignments were first “grouped by chromosome” to quickly visualize where short-read paired mates map to other chromosomes. It was expected that this grouping would align with ALT chromosomes identified *as BND calls with* Delly. Each integration site had an increase in short-read coverage for the 4 bp to 5bp that is duplicated around the LVV inserted sequence during repair of the staggered cuts. While paired-end short-read alignments were useful for identifying integration sites, they *cannot generally confirm the* expected full integration of the LVV sequence. To confirm full integration PacBio long-read alignments and variant calls were used. To cover all possible integrations, VCN2, VCN3 and VCN5 were sequenced with PacBio. It is expected, given a diploid genome, that all reads from one haplotype will support the integration at a given site. For PacBio, BWA alignments generally have split reads at integration sites, where part of the read maps to the GRCh38 reference chromosome and the remainder to the LVV chromosome as a supplementary alignment. In contrast, pbmm2 has more reads aligning across the integration site with an insertion that corresponds to the LVV inserted sequence, with no supplementary alignments to the LVV chromosome if the read contains the insertion. Pbsv insertion variant calls at the pbmm2 PacBio alignment integration sites were used to confirm complete insertion of the LVV sequence. To confirm integration of the LVV sequence and that no partial integrations were missed, all pbsv identified insertions greater than 50 bp were aligned back to the modified reference to determine if they aligned to the LVV sequence. These alignments were visualized in IGV. All previously identified integrations were confirmed and no additional integrations were present.

Single nucleotide variants (SNVs) and a 1 bp deletion were identified within the integrated sequences when pbsv insertion calls aligned to the LVV chromosome in the modified reference. We evaluated support for the putative variants in PacBio and Illumina reads. Generally, if a variant was present, the fraction of reads supporting the variant should be consistent with the fraction of insertions within the genome containing the variant.

## Results

### Stable clonal cell lines with 1 to 5 copies of integrated CD19 CAR

CD19 CAR lentivirus transduced Jurkat cells were selected with 10 μg/mL of Bleomycin for the first 3 passages and then with 5 μg/mL of Bleomycin. The Bleomycin resistant cells were subjected to single cell clone printing and screening. The single cell clones were screened by ddPCR using RRE assay and S2 as single copy control gene. Initial transduction using MOI = 1 and 5 resulted mostly VCN1 clonal cell lines. We only obtained two VCN2 clonal cell lines with no GFP expression. We chose one of the VCN1 clonal cell lines for lentiviral transduction to generate higher VCN clonal cell lines (as shown in Fig. 1B). Finally, we used VCN4 clonal cell line to generate VCN5 clonal cell line (Fig. 1B). We analyzed the cells by flow cytometry to detect the GFP and CD19 CAR expression level. The first-round screening resulted in many clones that either lost GFP expression after a few passages or CD19 CAR expression was not correlated with the integrated VCN (data not shown). Thus, we did a second lentivirus transduction on the selected VCN1 clone based on GFP expression. With the same screening process, we obtained VCN2, VCN3, and VCN4 clones that had GFP and CD19 CAR expression correlated with the integrated VCN. We further transduced VCN4 with the same lentivirus vector to obtain VCN5 with one additional integration. The GFP expression levels were similar between all the cell lines (Fig. 2A), however the CD19 CAR expression levels were correlated with the integrated VCN (Fig. 2B and 2C) except VCN1 cells, which exhibited GFP positive but no detectable CD19 CAR expression.

**Fig. 2.**
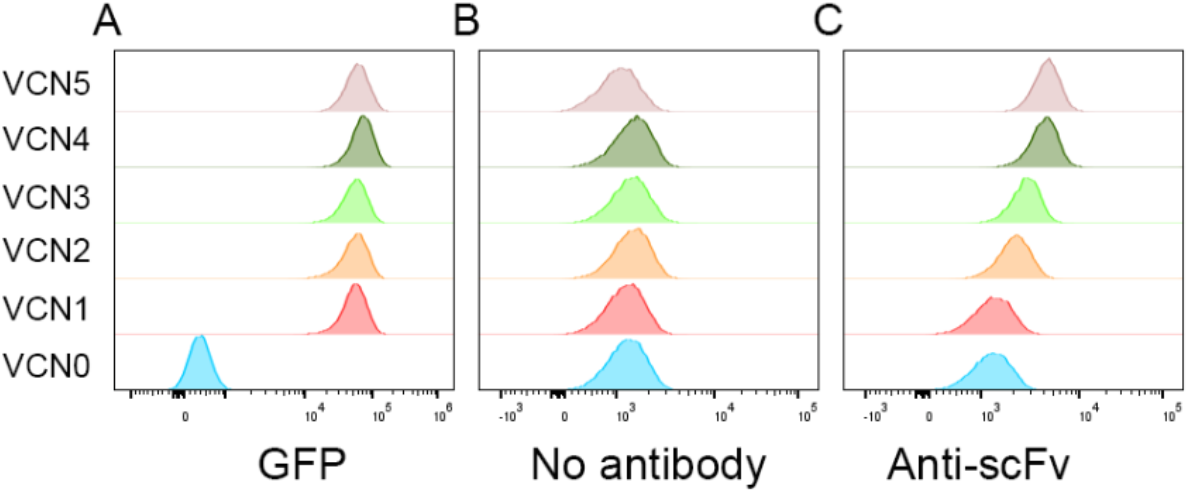
GFP and CD19 CAR expression in VCN clonal cell lines. A, GFP expression of each clonal cell lines analyzed by flow cytometry. B, No primary antibody controls for analyzing CD19 CAR expression by flow cytometry. C, CD19 CAR surface expression on each clonal cell line. Data are representative of 3 independent experiments.

### Measurement of VCN by different PCR targets

We utilized the assays developed based on the lentivirus elements (WPRE, RRE, Psi and LTR assays were published previously)^35^ and the CD19 CAR. S2 and RPL32 (Ribosomal protein L32) ^41^ were used as single copy reference genes. Most of the assays resulted in the expected copy number for each VCN cell line (Fig. 3). However, VCN3 appeared to have additional copy for WPRE, and both VCN4 and VCN5 appeared to have an extra copy of eGFP, results corroborated below by long-read NGS.

**Fig. 3.**
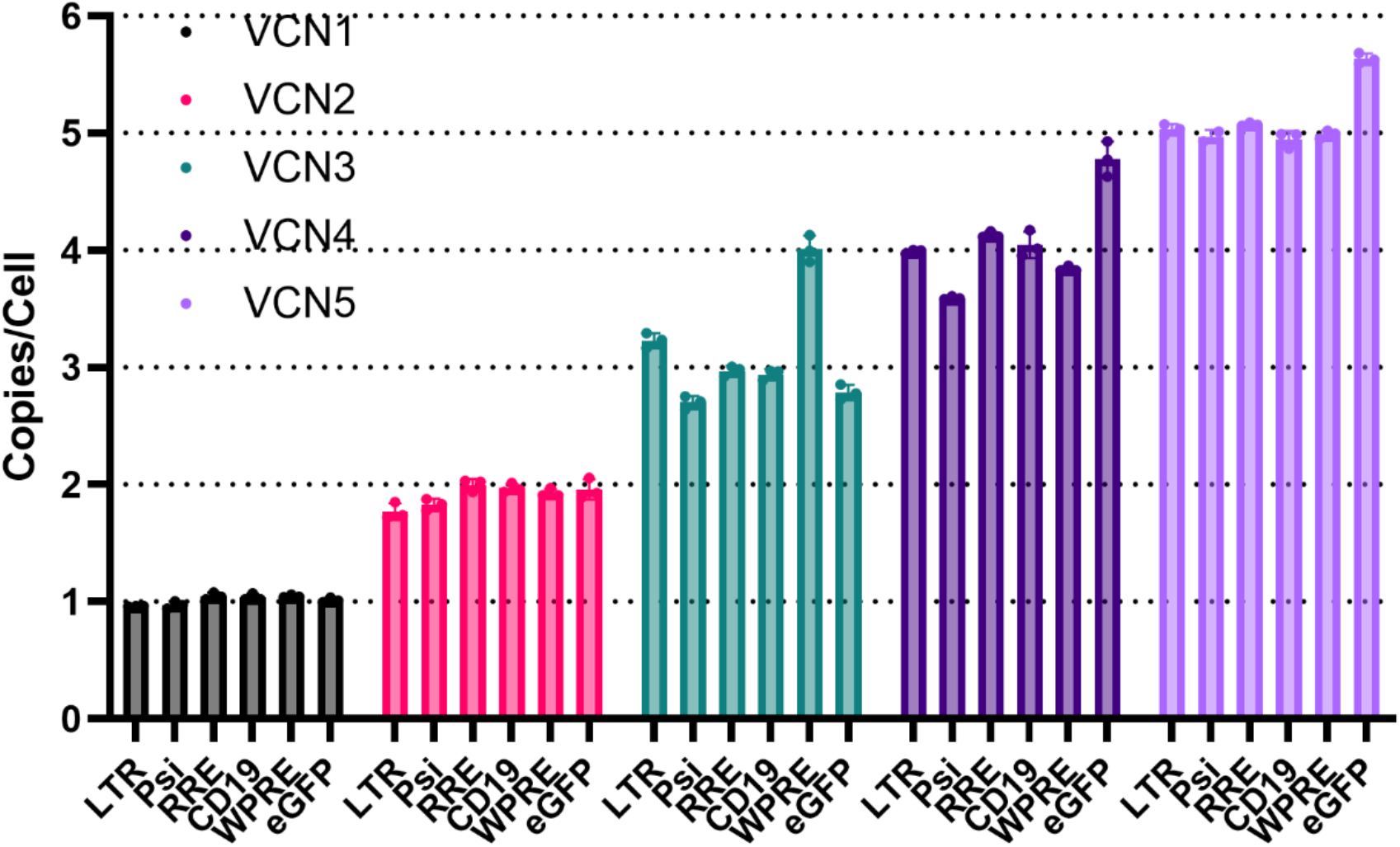
VCN detected by 6 different assays in each clonal cell line. Extra copy of WPRE was detected in VCN3 cell line. An extra copy of eGFP was detected in VCN4 and VCN5 cells. Representative of 3 experiments. Error bars represent standard deviation of the mean (n=3). Y-axis: Vector copy number / (copies/cell).

### Stability of the clonal cell lines

The stability of the cell lines was assayed by passaging the cells up to passage 42. Genomic DNA was extracted from the cells every week (3 passages per week) for 8 weeks and was analyzed by ddPCR to ensure the VCNs based on RRE and CD19 CAR. As shown in Fig.4, the VCN cell lines exhibited minimal variations for at least eight weeks, which indicated that the cell lines remained stable for the VCN measurements.

**Fig. 4.**
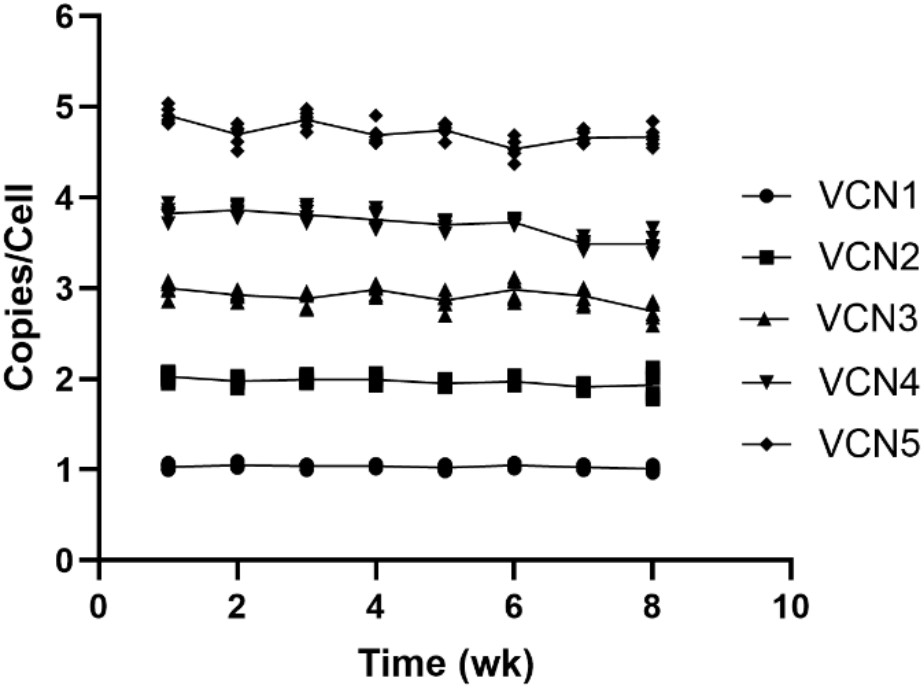
Stability of clonal cell lines. VCN of each cell line was determined based on ddPCR results of RRE and CD19 CAR. Data are representative of three independent experiments, each performed in technical triplicate for both RRE and CD19 CAR assays (n = 6 point per experiment). Error bars represent the standard error of the mean (SEM). X-axis: Time / weeks.

### Long-read sequencing identifies structural and small variants in provirus integrations

A combination of long-read and short-read whole genome sequencing identified and confirmed precise integration sites, target site duplicated sequences, and variation within each inserted sequence in the Jurkat cell genome (Fig. 5B). All insertions occurred in introns except for one in the 3’ UTR of *HNRNPR*. Interestingly, of the 10 unique integration events, only 3 contained the expected sequence, with others containing substitutions, deletions, and/or duplications relative to the expected sequence. VCN1 clonal cell line showed one copy of proviral DNA inserted in minus strand of chromosome 17 at position of 78 730 345. Long-read sequencing revealed a 724 bp deletion in the EF1α promoter region (Fig. 5B), which explained why VCN1 cells did not express CD19 CAR (Fig. 2C). This deletion is not visible by short-read sequencing due to low mapping quality caused by high identity with the EF1α promoter in the reference. Since VCN2-5 clonal cell lines were all derived from VCN1, all the cell lines had a copy of 724 bp-deletion EF1α promoter proviral DNA on chromosome 17 (Fig. 5B). As expected, VCN2 cells contained an additional insertion. However, it harbored four unexpected substitutions. Although both short-reads and long-reads detected these substitutions, only long-read data provided sufficient context to confirm that all four were located within the second insertion (Fig. 5, yellow lines). In VCN3 cells, one of the new proviral insertions had the precise expected sequence. However, the other proviral insertion had a mostly as expected sequence except one SNV, and an additional 802 bp sequence on the 3’ end of proviral insertion. This additional sequence contains a 778 bp duplication of a portion of the WPRE sequence and 24 bp of inverted 3’ LTR sequence. This 778 bp duplication accounts for the additional WPRE copy detected by the WPRE ddPCR assay in the VCN3 cell line (Fig. 3). Interestingly, both short- and long-read whole genome sequencing data revealed an additional non-functional 258 bp insertion of predominantly LTR sequence that was not detected by ddPCR assay. Analysis of VCN4 revealed one new proviral insertion exhibiting the expected sequence, a second containing a single nucleotide substitution, and a third harboring both with a substitution and additional 41 bp of sequence at downstream of the 3’ LTR. This additional 41 bp sequence comprises 15 bp of unknown origin and 26 bp derived from 3’ LTR (Fig. 5B). Furthermore, short- and long-read sequencing data identified an additional 2070 bp integration corresponding to the 3’ region of the proviral DNA. This fragment contained a 1 bp deletion and encompassed the eGFP gene, thereby accounting for the extra copy of eGFP detected by ddPCR assay in the VCN4 cells (Fig. 3). The VCN5 cell line contained all the insertions previously identified in VCN4, alongside one additional insertion exhibiting the precise expected sequence (a representative visualization utilized for manual curation is provided in Supplemental Fig. S1). Furthermore, the allele frequencies of SNVs and insertions/deletions (indels) within the LVV sequences, as determined by both short- and long-read sequencing, corresponded with the expected fraction of variant-containing copies (Supplemental Table S1). All reported integrated sequences derive from the HiFi pbmm2-pbsv calls, starting with the 4 bp to 5 bp of duplicated integration site sequence, and are available in Supplementary File 1. In summary, whole-genome sequencing revealed alterations within multiple inserted sequences, thereby elucidating the unexpected PCR results.

**Fig. 5.**
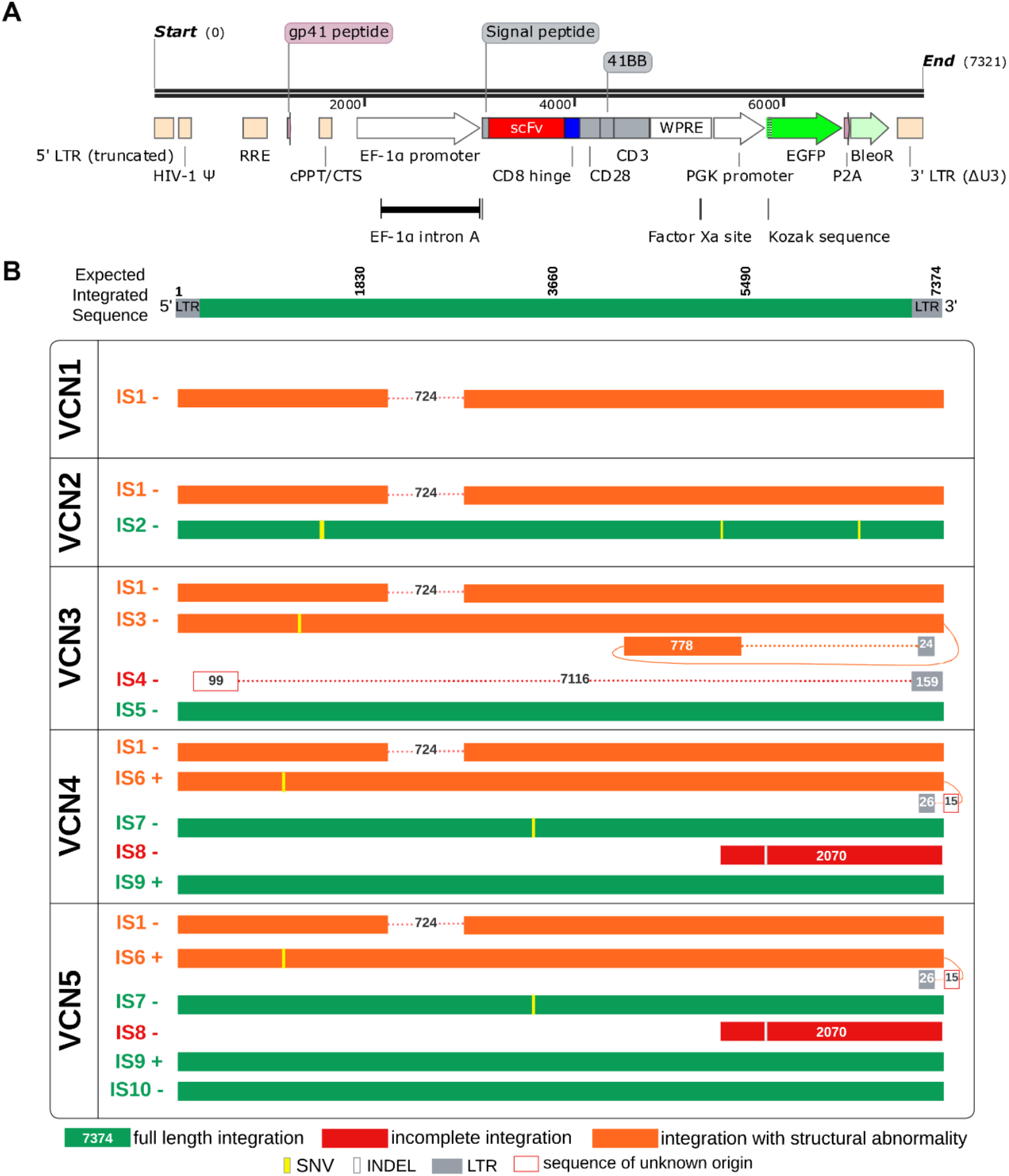
Schematic diagram of lentiviral DNA insertions and variants inside the insertions in each clonal cell line. A. schematic diagram of each element of the lentiviral DNA insertion. The truncated 5’ LTR is replaced by the 3’ LTR during integration. B. Schematic diagram of proviral DNA integration sites across VCN1 - VCN5 characterized using short- and long-read sequencing. Integrations are categorized as full-length (green), structurally abnormal (orange), or incomplete (red), with genomic orientation indicated by +/- symbols. Some integrations occur in multiple VCNs, *e*.*g*., IS1 with a 724 bp deletion in EF1a promoter occurs in all clonal cell lines because higher copy number clones were developed from VCN1. Horizontal dotted lines signify missing regions, while vertical yellow and white lines denote SNVs and INDELs, respectively. Grey boxes represent LTR sequence. Curved orange lines connect unexpected sequence post expected sequence. Some unexpected sequences of unknown origin were observed and noted by a red outlined box. More information about each integration is in Table 1 and Supplemental Table S2.

Analysis of both short-read and long-read whole genome sequencing data also revealed the absence of chromosome Y in VCN1 to VCN5 cell lines. A previous report ^42^ and our G-band karyotype analysis (Supplemental Fig. S2A) indicated that a subpopulation of Jurkat cells lacks chromosome Y (7 out of 20 metaphase spreads). In this work, sequencing and G-band karyotyping confirmed this absence of chromosome Y in the VCN1 cell line (Supplemental Fig. S2B). Because all the other VCN cell lines were derived directly from VCN1, this shared lineage accounts for the absence of chromosome Y across all the VCN1 to VCN5 clonal cell lines.

**Table 1.**
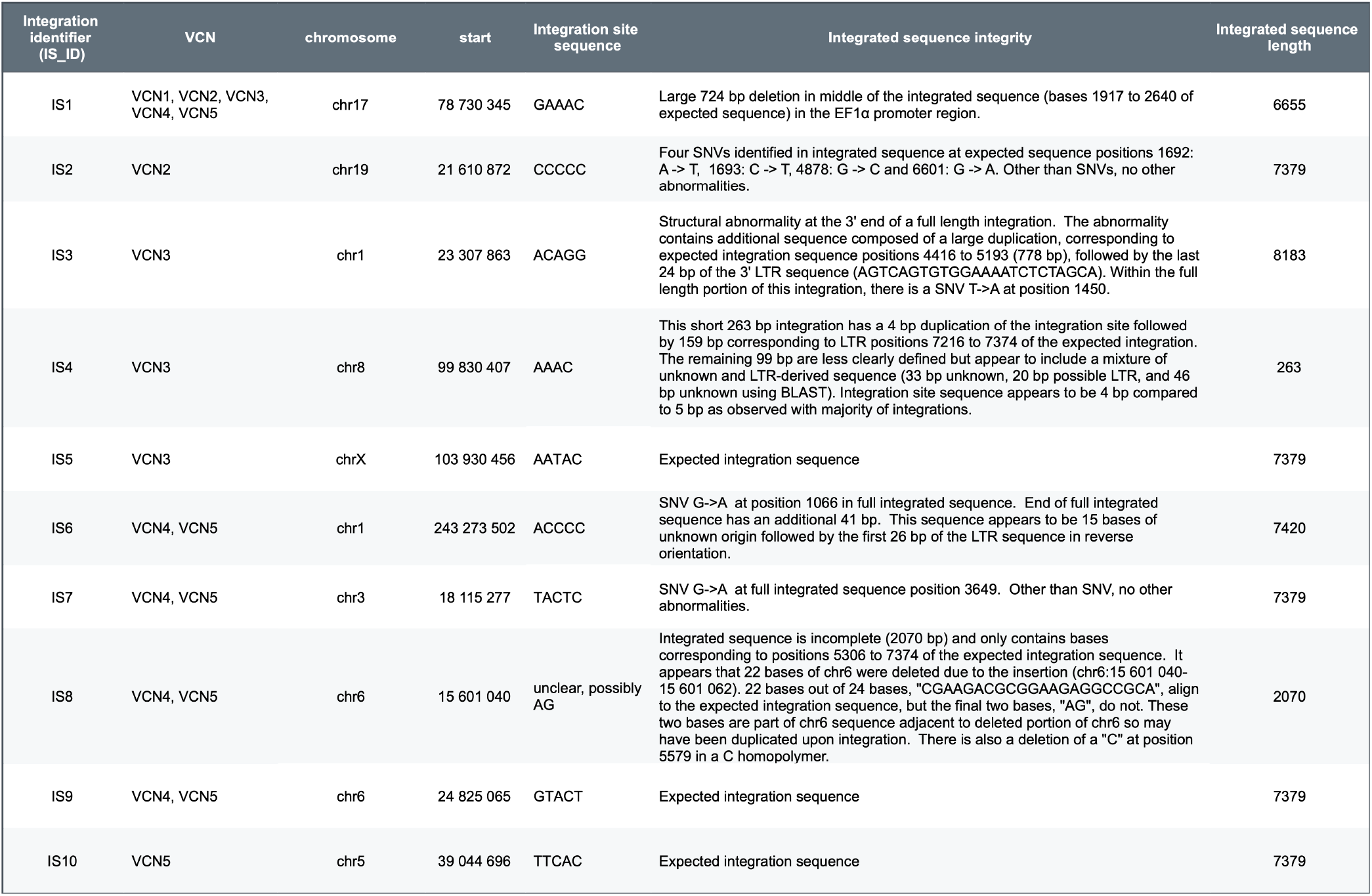
Summary of all integration sites and integration observations. The IS_ID corresponds to chromosome integration site; IS_IDs can be shared across VCN. Integration positions on the chromosome (start/end) are the first and last base of the integration site, they are not left aligned. Positions of abnormalities are relative to the expected proviral sequence, not the position within the sequence reported in the fasta format. A full summary of all integrations and visualization of integration sites are listed in Supplemental Table S1 and Supplemental Fig. S3, respectively.

## Discussion and conclusions

In this study, we generated deeply characterized clonal cell lines as reference samples for LVV integration site, quality, and copy number. With short-read and long-read whole genome sequencing, we identified all LVV integration sites and sequences, including some small segments of the LVV in the cell lines (Fig. 5)^38^. Long-read WGS uncovered the structural complexity of LVV integrations that were invisible to standard short-read methods. While short-read sequencing is effective for identifying general integration sites, it often fails to resolve full insertion sequences or detect large internal rearrangements due to low mapping quality in repetitive or high-identity regions and inability to resolve full insertions. A primary example in this study is the characterization of all the integration sites present in the clonal cell lines, where long-read sequencing revealed a 724 bp deletion in the EF1α promoter region. This specific deletion explained why the VCN1 cells failed to express the CD19 CAR protein despite the presence of the proviral DNA. This was invisible to short-read sequencing because the deletion occurred in a region with high identity to the native EF1α promoter in the human reference genome. Beyond simple deletions, long-read data identified complex variants across the clones, revealing that only 3 out of 10 unique integration events contained the precise expected sequence. The remaining integrations featured unexpected substitutions, deletions, and duplications, such as the 778 bp duplication found in a VCN3 insertion and a 2070 bp partial insertion in VCN4 that included an extra copy of the eGFP gene. These findings highlight that LVV integration is not always a clean, full-length process and that long-read sequencing is essential for the high-resolution characterization required to validate reference materials and may help ensure the safety of CAR-T cell therapies.

To echo the findings in these CAR VCN materials, we also found SNVs in the proviral DNA regions in VCN4, VCN2 and VCN1 samples of NIST Research Grade Test Material (RGTM) 10268 ^43^, and a 648 kb genomic deletion near the integration site chr14:22564447 in the VCN3 sample (unpublished data from the RGTM 10268) ^44^.

There are some limitations of this study. First, we generated our clonal VCN cell lines using Jurkat cells, a leukemia cell line known for inherent genomic instability, including frequent mutations, chromosome rearrangements, and single nucleotide polymorphism^45^. Because of this instability, it is possible that the variant lentiviral DNA insertions we observed were a byproduct of the host cells’ tendency to rearrange or truncate DNA during or after integration. However, we did not observe partial lentiviral DNA insertions in our previous study ^32^, which indicated that the likelihood of Jurkat cell genome causing these specific partial insertions is low. In addition, a new targeted long read approach identified similar promoter deletions in a variety of non-clonal CAR-T products ^29^. In the future, since we established a method for immortalizing human primary T cells ^46^, we plan to collect explicitly consented patient primary cells to establish immortalized T cell lines, which would minimize the possibility of rearrangements or mutations in the cell line. Second, this set of materials have yet to be validated by multiple laboratories to prove its broad utility and commutability.

In conclusion, we developed a set of VCN clonal cell lines as candidate reference samples, aimed to assure confidence in measurements of one to five LVV copies, as well as integration sites and sequences to evaluate insertion quality.

## Abbreviations

CAR: Chimeric antigen receptor
VCN: Vector copy number
LTR: Long terminal repeat
ddPCR: Droplet digital polymerase chain reaction
eGFP: Enhanced green fluorescent protein
FDA: Food and Drug Administration
ALL: Acute lymphoid leukemia
CRISPR: Cluttered regularly interspaced short palindromic repeats
LVV: Lentiviral vector
NGS: Next generation sequencing
RGTM: Research grade test material
WHO: World Health Organization
WPRE: Wood chuck Hepatitis Virus Postranscriptional Regulatory Element
PGK: Phosphoglycerate Kinase
MOI: Multiplicity of infection
RPL32: Ribosomal protein L32
IS: Integration sites
SV: Structural variant
SNV: Single nucleotide variant
Indels: Insertion and deletions
WGS: Whole genome sequencing

## Disclaimers

Certain commercial equipment, instruments, and materials are identified to specify the experimental procedure. In no case does such identification imply recommendation or endorsement by the National Institute of Standards and Technology, nor does it imply that the materials or equipment are necessarily the best available for the purpose.

## Author Contributions

Conceptualization, ZH, JHM, LW, SL-G, JMZ, H-JH; methodology, ZH, JMH, LT, MM,NX, SL-G, JMZ, H-JH; investigation, ZH, JHM, LT, MM, NX; data curation, ZH, JHM, LT, JMZ; resources, ZH, LW, SL-G, JMZ,H-JH; writing-original drift preparation, ZH, JHM, JMZ; writing-review and editing, ZH, JHM, LT, MM, NX, LW, SL-G, JMZ, H-JH; visualization, ZH, JHM, LT, NX.

## Data Availability Statement

Whole genome sequencing data is available on public databases (Bioproject Accession: [PRJNA1282768](https://www.ncbi.nlm.nih.gov/bioproject/1282768))

## Consent for publication

The National Institute of Standards and Technology Research Protections Office reviewed the protocol for this project and determined it is “not human subjects research” as defined in 15CFR27, the Common Rule for the Protection of Human Subjects. The Jurkat cell line was derived from tissues from a 14-year-old with acute T-cell leukemia in the 1970s. The cell line was established prior to the current research use consent process. In 2013, The Cancer Genome Atlas (TCGA) steering committee administered by NCI and NHGRI at NIH decided that the benefits outweighed the risks for making genomic data fully public without controlled access for Jurkat and other widely-used, commercially-availably cell lines in the Cancer Cell Line Encyclopedia https://www.cancer.gov/ccg/research/structural-genomics/tcga/history/policies/ccle-open-release-justification.pdf. For the Jurkat cell line, short-read whole genome sequencing data was previously published (https://bmcgenomics.biomedcentral.com/articles/10.1186/s12864-018-4718-6) without controlled access in public databases, so there is minimal additional risk of reidentification for publishing the long-read whole genome sequencing data in this manuscript. To make progress towards similar cell lines with explicit consent for public genome data sharing and cell line development, NIST is currently working towards establishing T cell lines and other types of cell lines from individuals who have provided consent under the Personal Genome Project (https://www.pnas.org/doi/abs/10.1073/pnas.1201904109). This consent includes permission to share genomic data and cell lines publicly for open research and commercial use, understanding the risks associated with this. For these reasons, the NIST Research Protections Office determined that the whole genome sequencing data generated in this work could be made publicly available.

## Acknowledgments

We thank Dr Jerilyn R. Izac and Dr. Simona Patange for their critical reading, insightful comments, and valuable suggestions.

