## Supplemental Figs S1-S3, Table S1 for "Long-read genome sequencing reveals complex variability in lentiviral provirus insertions in deeply characterized Clonal CD19 CAR-T vector copy number reference cell lines"

**Supplementary Information**

1. Supplemental Fig. S1
2. Supplemental Table S1
3. Supplemental Fig. S2
4. Supplemental Fig. S3

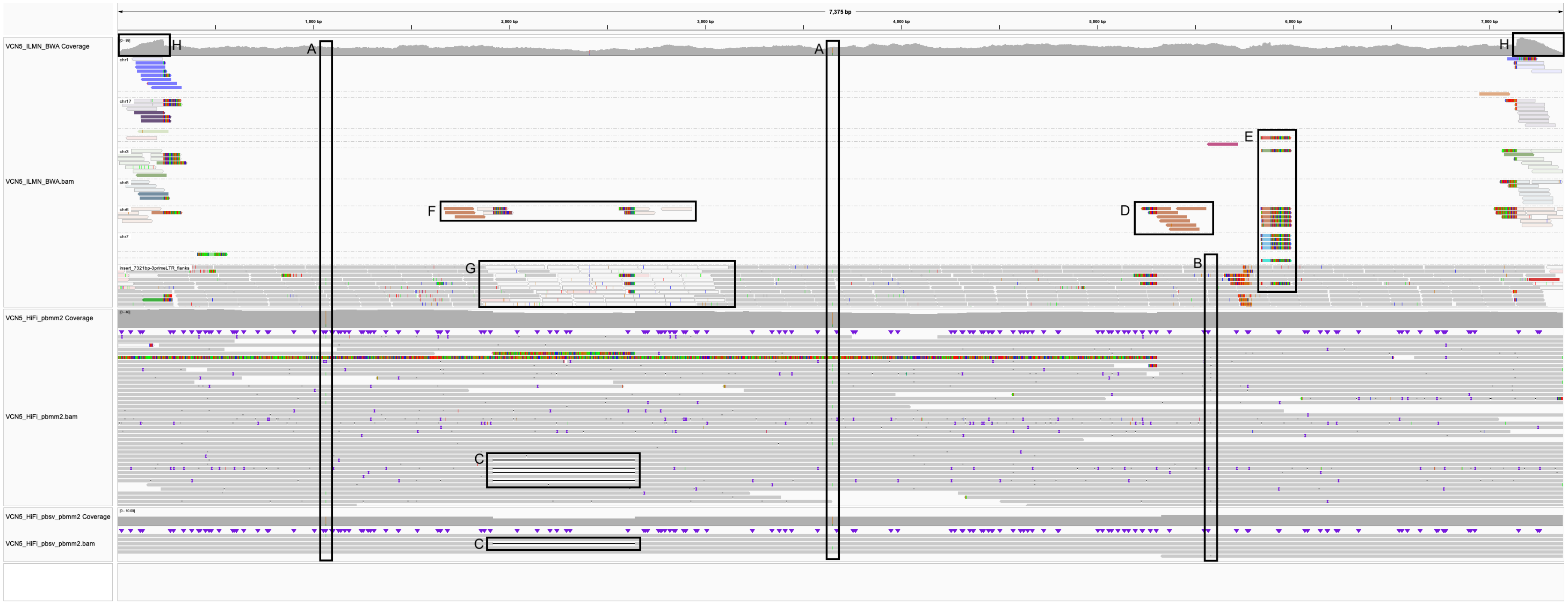


**Supplemental Fig. S1 Alignments of Illumina reads, HiFi reads, and pbsv variant calls to the provirus sequence for VCN5.**

The Illumina track is grouped by ”chromosome of mate” to highlight integration sites. Visualization with IGV identified specific variants and mapping artifacts annotated as follows: A) SNVs observed on integrated sequence in chr1 and chr3. Note, IGV "coverage allele fraction threshold" = 0.15. B) DEL observed on integrated sequence in chr6. Given the extra insertion (IS8) in VCN5, it was expected that the DEL would be observed in 17% of reads. While not visible in the screen shot, 10 % (6/58) of Illumina reads contained the DEL. The DEL can be seen in PacBio reads and was observed in 19 % (6/32) of reads. With PacBio reads it was possible to see supplementary alignments of the reads supporting the deletion, with four correctly on chr6 but two from other chromosomes, likely due to indel errors in PacBio. C) 724 bp deletion of promoter in integration sequence on chr17. D) Partial insertion (2046 bp) on chr6. E) VCN5, but not other VCNs, have Illumina reads partially aligned to ~60bp, but this is not seen in HiFi reads in any VCN. F) ILMN read mates map incorrectly with MQ0 to the *EEF1A1* (chr6:73.49-73.53) gene in GRCh38 because this human gene promoter was used in the provirus. This was observed with all VCN ILMN reads. G) Region has homology with *EEF1A1* gene in GRCh38, causing zero mapping quality in ILMN data but not long reads. This was observed with all VCN ILMN reads. H) Flanking LTR regions have high coverage of ILMN reads with low MQ because these sequences are identical.

| | VCN2 | VCN2 | VCN2 | VCN2 | VCN3 | VCN4/5 | VCN4/5 | VCN4/5 |
|---|---|---|---|---|---|---|---|---|
| **position in expected proviral sequence** | **1639** | **1640** | **4825** | **6548** | **1450** | **1066** | **3649** | **5579** |
| chromosome of inserted sequence with SNV/DEL in pbsv call | chr19 | chr19 | chr19 | chr19 | chr1 | chr1 | chr3 | chr1/5/6 |
| REF | A | C | G | G | T | G | G | C |
| ALT | T | T | C | A | A | A | A | DEL |
| **PACBIO PBMM2** | | | | | | | | |
| read support for ALT | 10 | 10 | 7 | 6 | 12 | 7 | 7 | 1/1/4 |
| total reads | 16 | 16 | 10 | 10 | 32 | 42 | 38 | 32 |
| **expect % support for ALT** | **50** | **50** | **50** | **50** | **33** | **20** | **20** | **17** |
| **observed % support for ALT** | **63** | **63** | **70** | **60** | **38** | **17** | **18** | **19** |
| **ILLUMINA BWA** | | | | | | | | |
| read support for ALT | 8 | 8 | 15 | 17 | 20 | 4 | 14 | 6 |
| total reads | 28 | 28 | 35 | 34 | 54 | 44 | 47 | 58 |
| **expect % support for ALT** | **50** | **50** | **50** | **50** | **33** | **20** | **20** | **17** |
| **observed % support for ALT** | **29** | **29** | **43** | **50** | **37** | **9** | **30** | **10** |

**Table S1 Support for the putative SNVs or deletion in PacBio and Illumina reads**.
Generally, if a variant was present, the fraction of reads supporting the variant should be consistent with the fraction of insertions in the VCN containing the variant. For example, if a SNV is in one of two insertions in a VCN clone, then we would expect approximately 50 % of reads for that integration site to contain a SNV at that position. For the deletion at 5579 the expected percent support for ALT is based on VCN5 since VCN4 was not sequenced with PacBio. Likely due to indel errors in the PacBio reads, this 2 reads supporting the deletion came from insertions on chr1 and chr5. SNVs all came from the expected chromosomes.

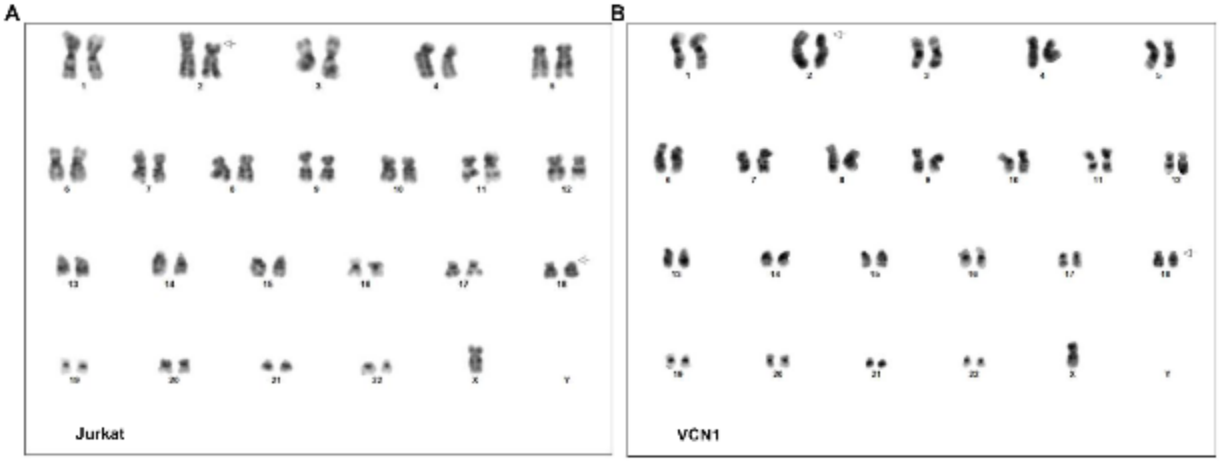


**Supplemental Fig. S2 G-band Karyograms of Jurkat cells and clonal cell line.**
A subpopulation of Jurkat cells and the clonal cell line VCN1 lacked the Y chromosome. A) G-band karyotype analysis revealed that a subpopulation (7 out of 20 metaphase spreads) of Jurkat cells lacked the Y chromosome. B) All VCN1 metaphase spreads exhibited a lack of the Y chromosome

**Supplemental Fig. S3 IGV screenshot of integration sites.**

Screenshots are grouped by genomic integration site for each VCN. Each series begins with the baseline control showing the integration site (Table 1) in the Jurkat genome without integration (VCN0). Where available, alignment tracks are provided for both HiFi reads (aligned with pbmm2) and Illumina reads (aligned with BWA-mem), corresponding to Fig. 1.

### IS1_VCN1-VCN2-VCN3-VCN4-VCN5_chr17:78,730,345

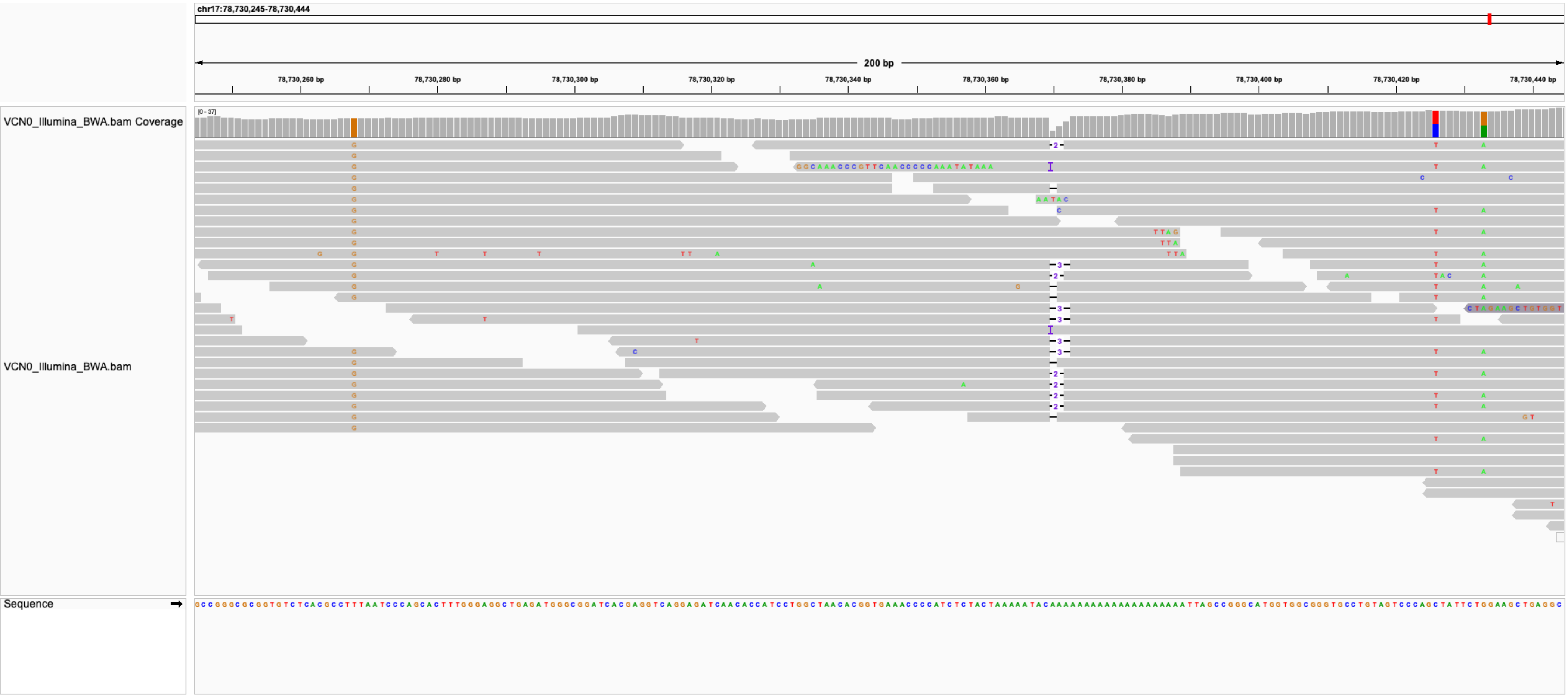

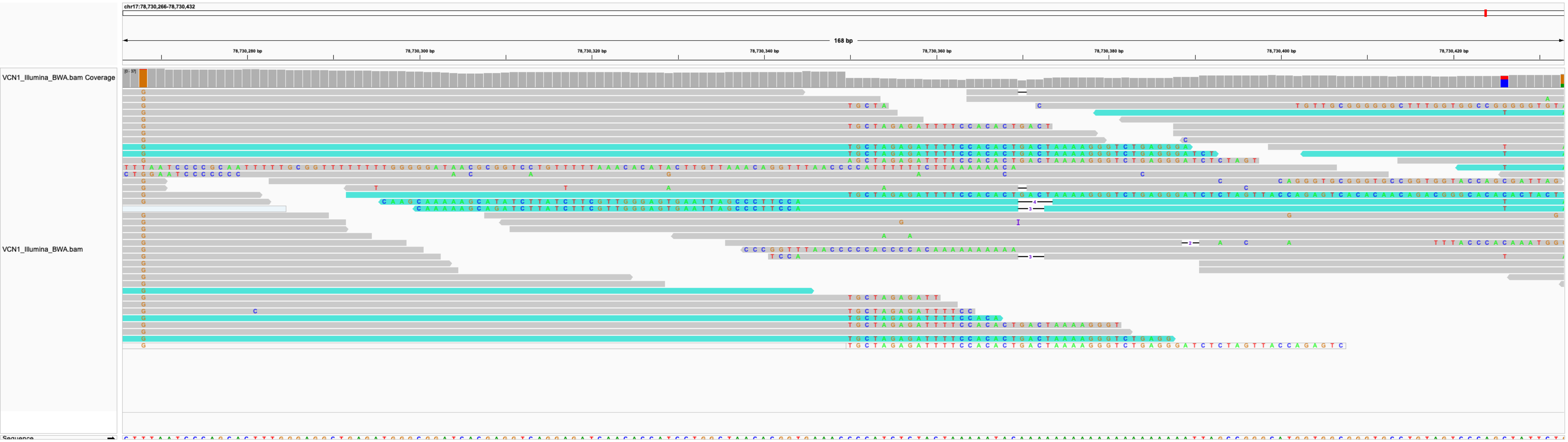
chr17:78,730,266-78,730,432
168 bp
VCN1_Illumina_BWA.bam Coverage
VCN1_Illumina_BWA.bam
Sequence

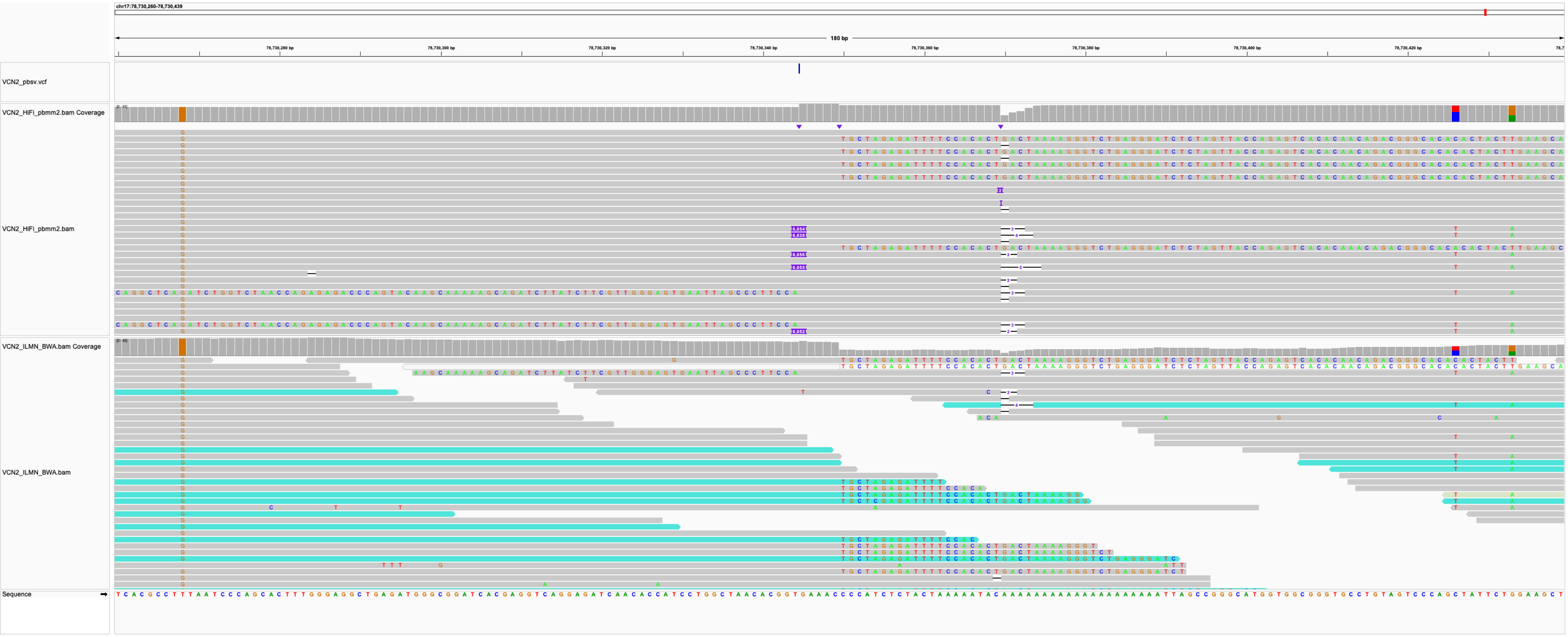
chr17:78,730,260-78,730,439
180 bp
VCN2_pbsv.vcf
VCN2_HiFi_pbmm2.bam Coverage
VCN2_HiFi_pbmm2.bam
VCN2_ILMN_BWA.bam Coverage
VCN2_ILMN_BWA.bam
Sequence

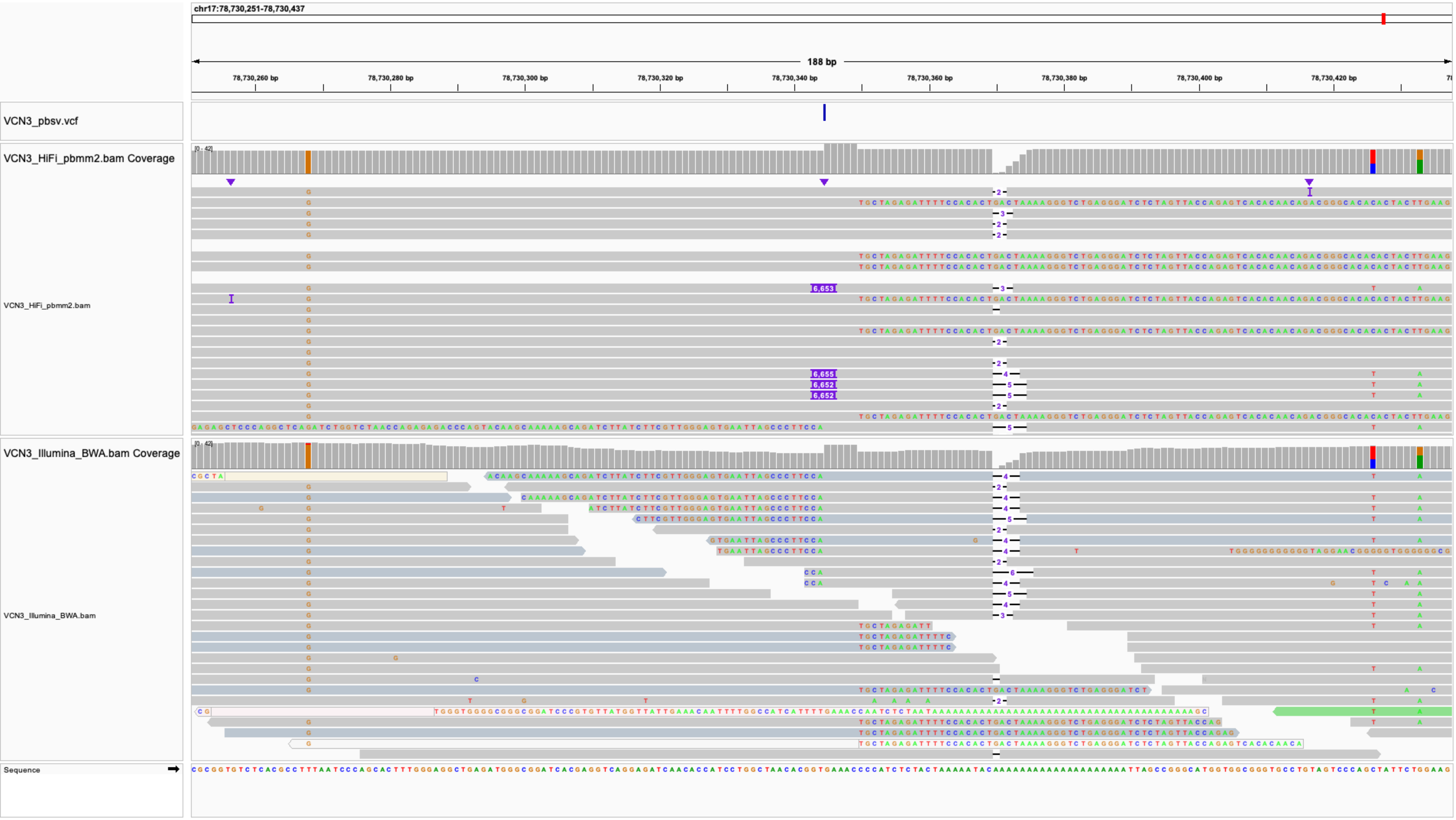
chr17:78,730,251-78,730,437
188 bp
78,730,260 bp
78,730,280 bp
78,730,300 bp
78,730,320 bp
78,730,340 bp
78,730,360 bp
78,730,380 bp
78,730,400 bp
78,730,420 bp
VCN3_pbsv.vcf
VCN3_HiFi_pbmm2.bam Coverage
[0 - 42]
VCN3_HiFi_pbmm2.bam
VCN3_Illumina_BWA.bam Coverage
[0 - 42]
VCN3_Illumina_BWA.bam
Sequence

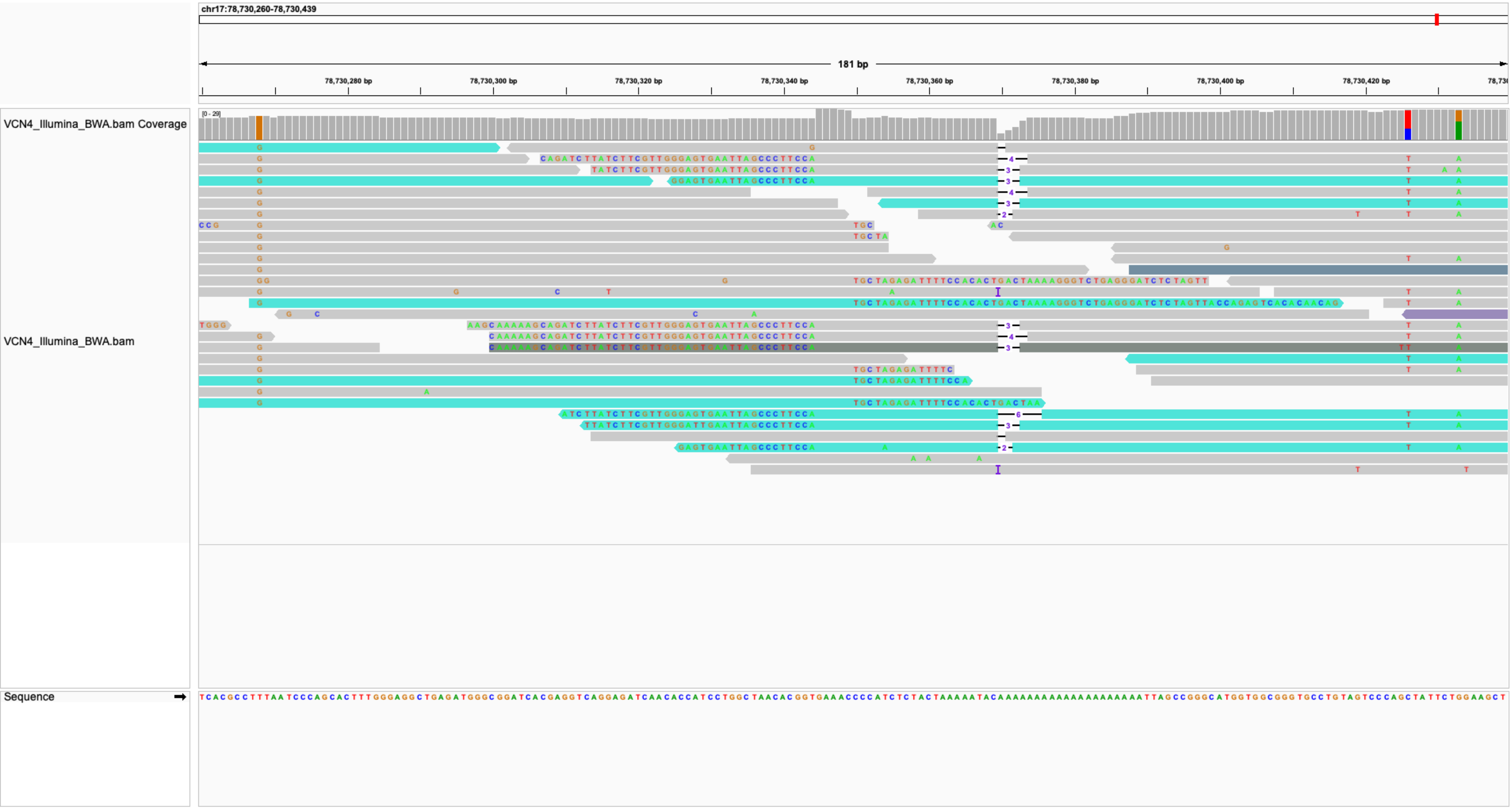
chr17:78,730,260-78,730,439
181 bp
78,730,280 bp
78,730,300 bp
78,730,320 bp
78,730,340 bp
78,730,360 bp
78,730,380 bp
78,730,400 bp
78,730,420 bp
VCN4_Illumina_BWA.bam Coverage
[0 - 29]
VCN4_Illumina_BWA.bam
Sequence

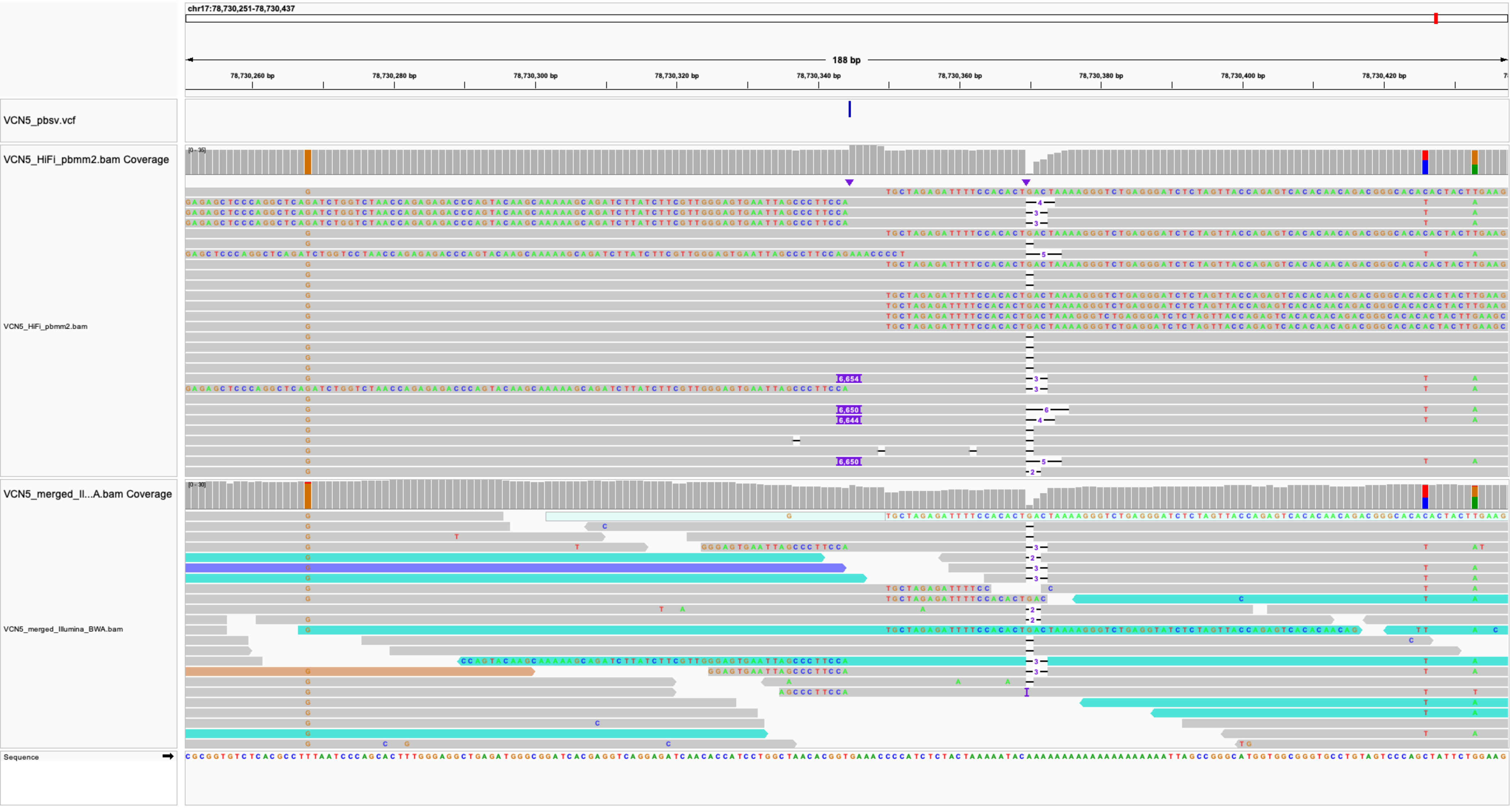
chr17:78,730,251-78,730,437
188 bp
78,730,260 bp
78,730,280 bp
78,730,300 bp
78,730,320 bp
78,730,340 bp
78,730,360 bp
78,730,380 bp
78,730,400 bp
78,730,420 bp
VCN5_pbsv.vcf
VCN5_HiFi_pbmm2.bam Coverage
VCN5_HiFi_pbmm2.bam
VCN5_merged_Il...A.bam Coverage
VCN5_merged_Illumina_BWA.bam
Sequence

### IS2_VCN2_chr19:21610872

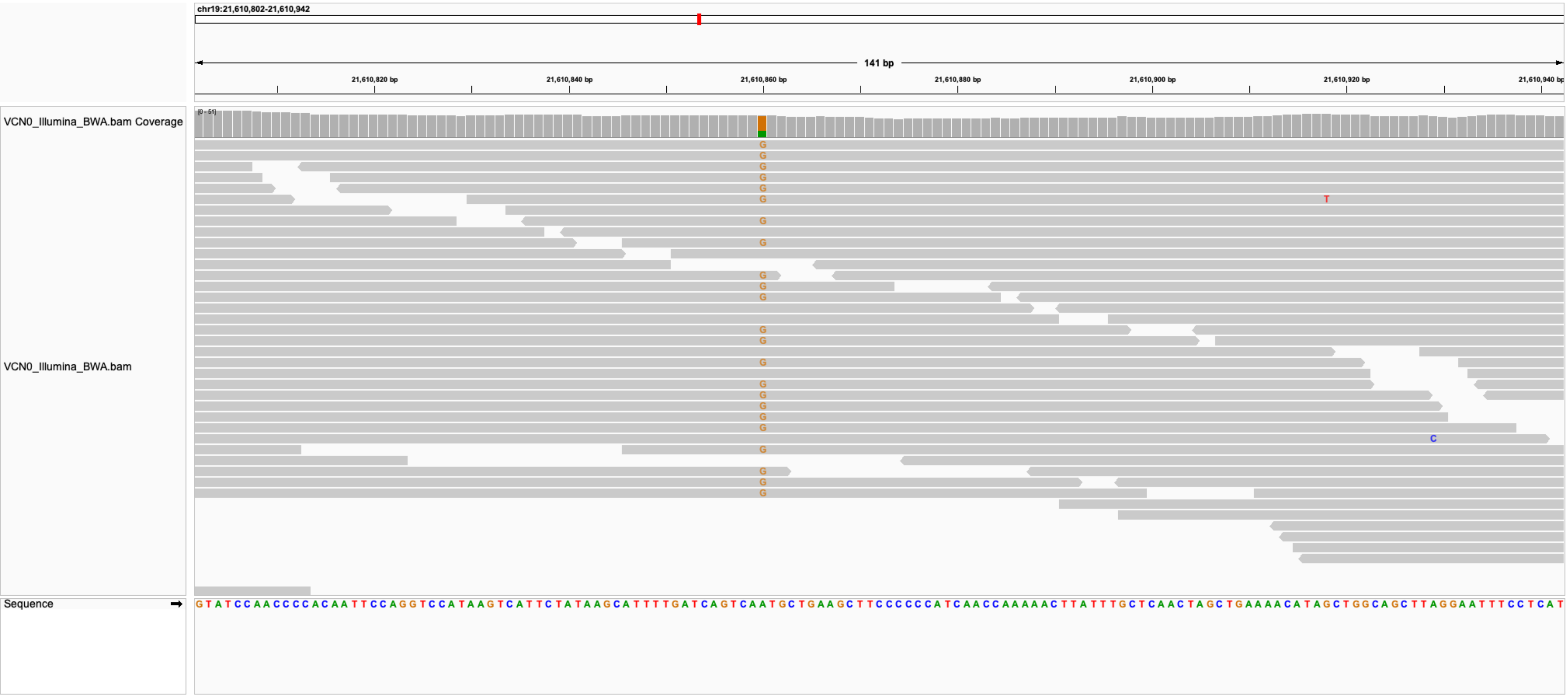

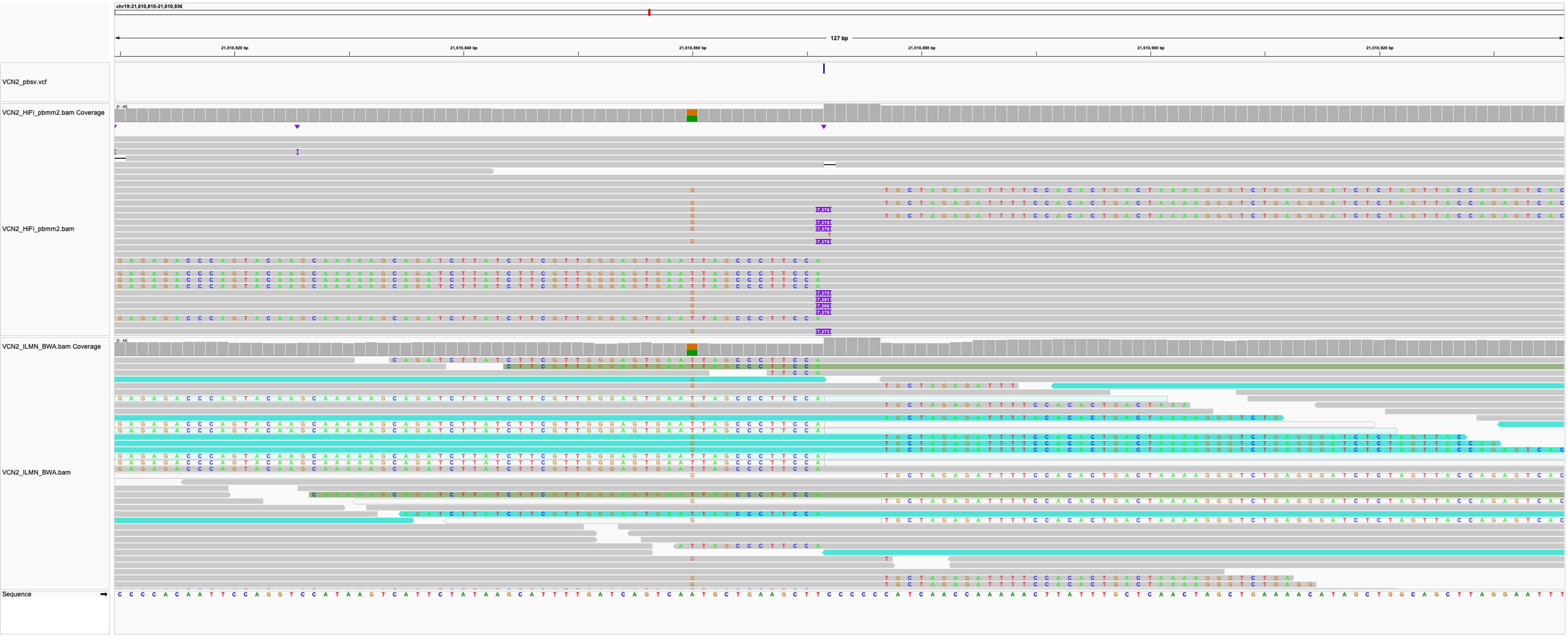

chr19:21,610,810-21,610,936
127 bp
21,610,820 bp
21,610,840 bp
21,610,860 bp
21,610,880 bp
21,610,900 bp
21,610,920 bp
VCN2_pbsv.vcf
VCN2_HiFi_pbmm2.bam Coverage
VCN2_HiFi_pbmm2.bam
VCN2_ILMN_BWA.bam Coverage
VCN2_ILMN_BWA.bam
Sequence

#### IS3_VCN3_chr1:23307863

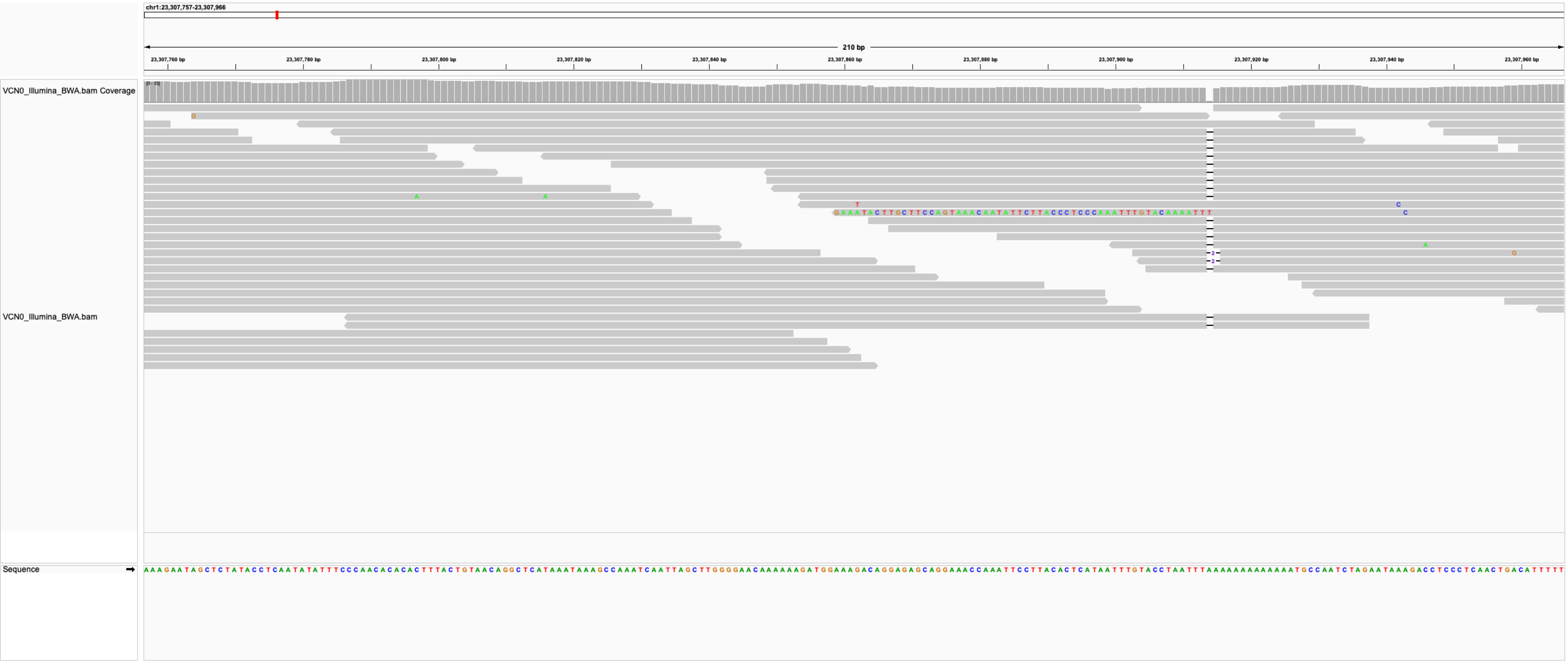

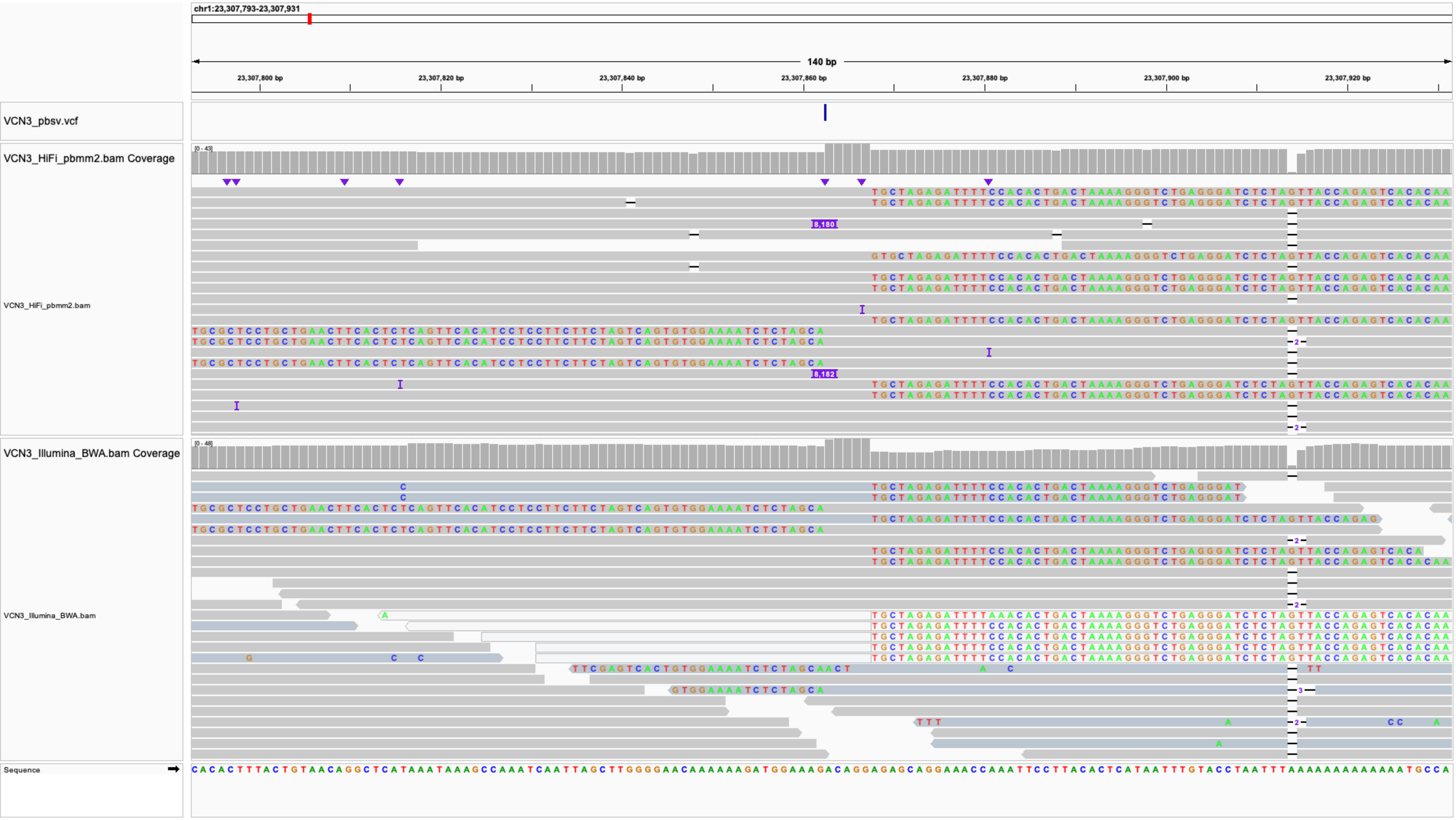
chr1:23,307,793-23,307,931
140 bp
23,307,800 bp
23,307,820 bp
23,307,840 bp
23,307,860 bp
23,307,880 bp
23,307,900 bp
23,307,920 bp
VCN3_pbsv.vcf
VCN3_HiFi_pbmm2.bam Coverage
VCN3_HiFi_pbmm2.bam
VCN3_Illumina_BWA.bam Coverage
VCN3_Illumina_BWA.bam
Sequence

### IS4_VCN3_chr8:99830407

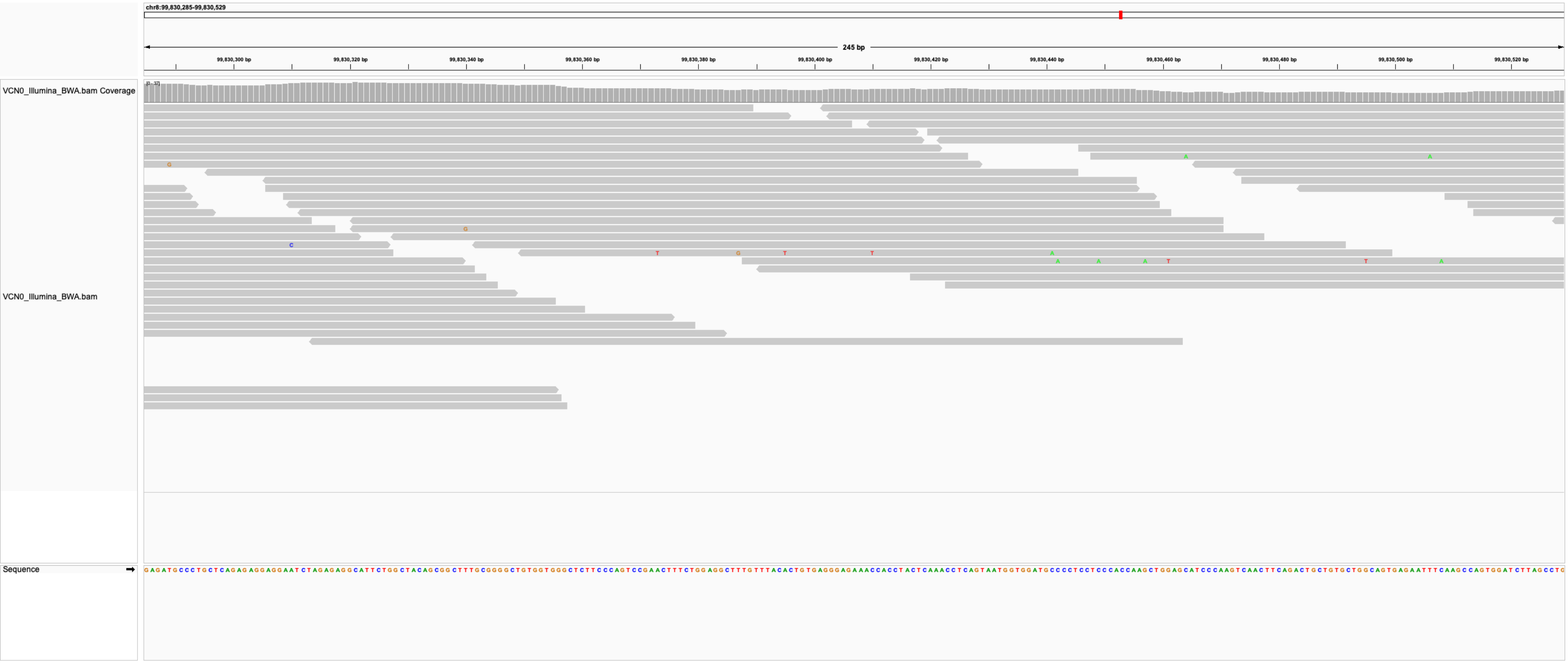

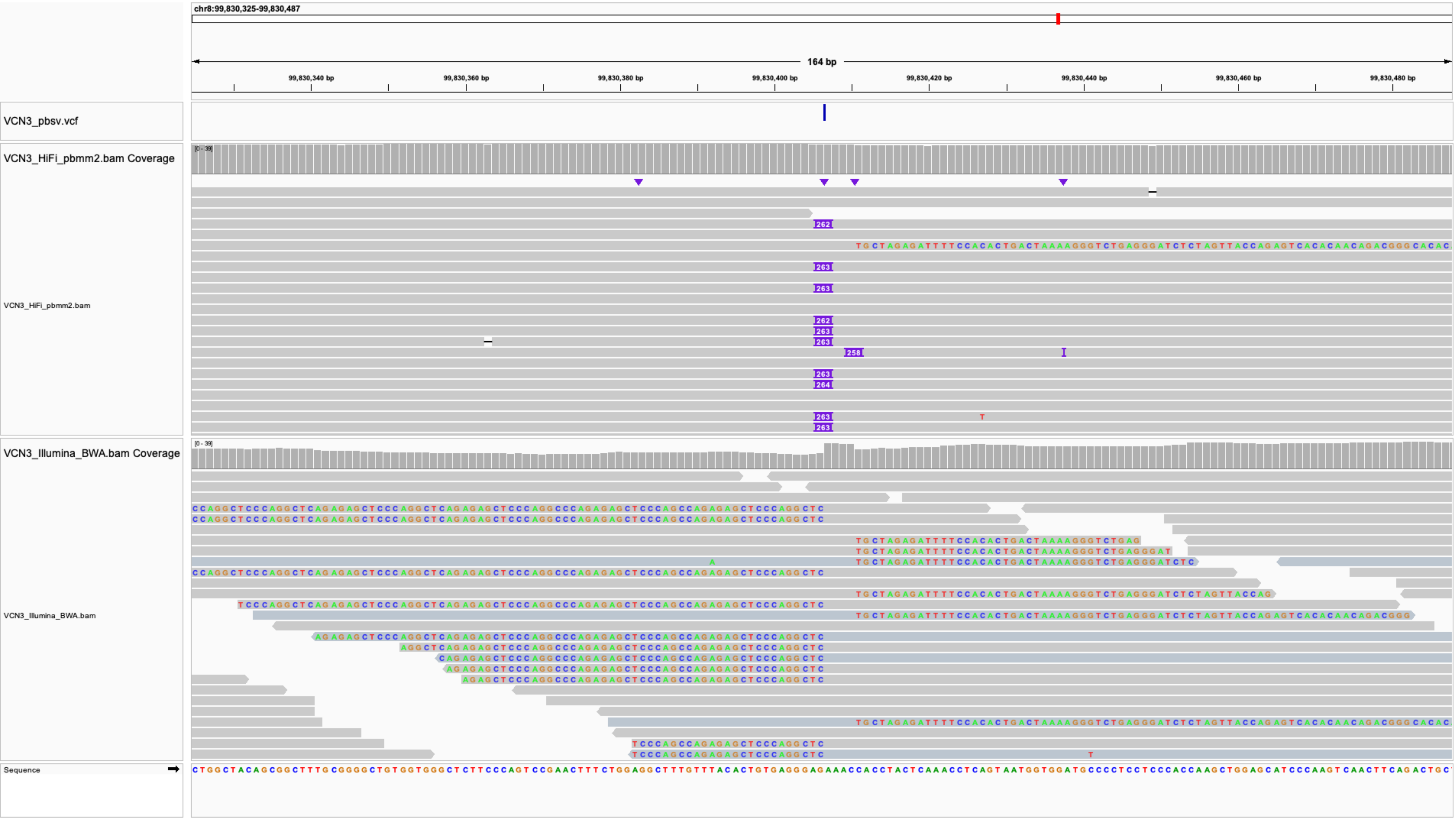

chr8:99,830,325-99,830,487
164 bp
99,830,340 bp
99,830,360 bp
99,830,380 bp
99,830,400 bp
99,830,420 bp
99,830,440 bp
99,830,460 bp
99,830,480 bp
VCN3_pbsv.vcf
VCN3_HiFi_pbmm2.bam Coverage
[0 - 39]
VCN3_HiFi_pbmm2.bam
262
263
263
262
263
263
258
263
264
263
263
VCN3_Illumina_BWA.bam Coverage
[0 - 39]
VCN3_Illumina_BWA.bam
Sequence

**IS5_VCN3_chrX:103930456**

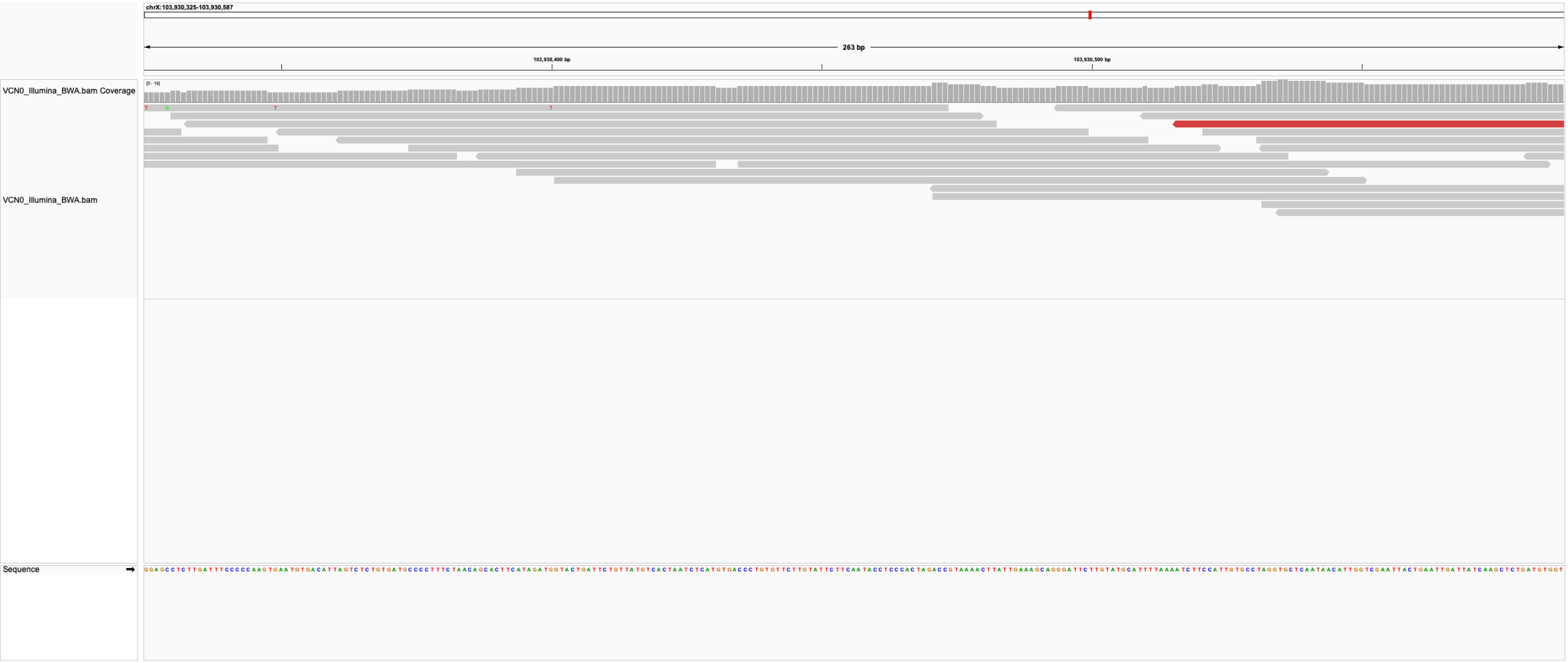

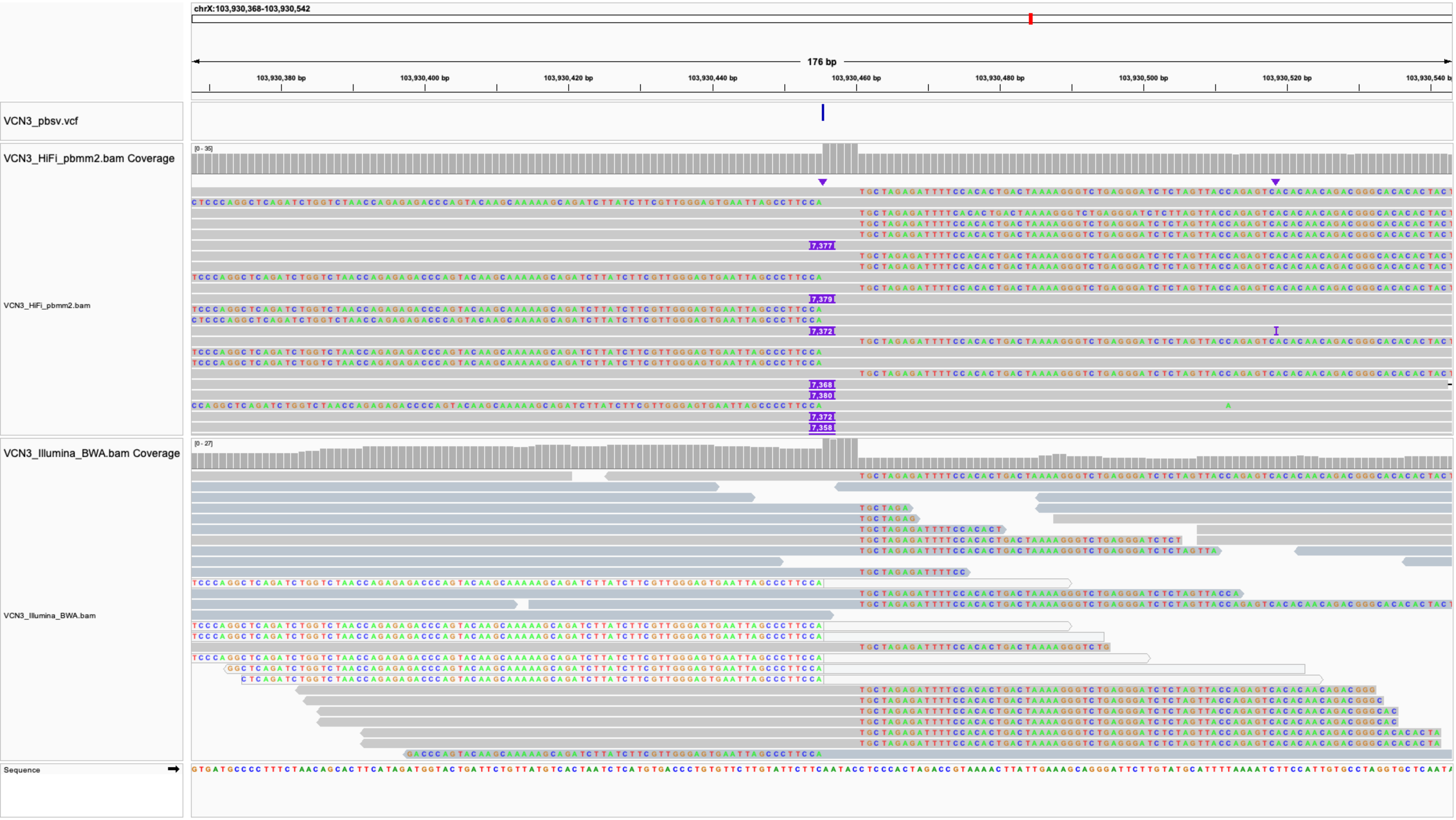
chrX:103,930,368-103,930,542
176 bp
103,930,380 bp
103,930,400 bp
103,930,420 bp
103,930,440 bp
103,930,460 bp
103,930,480 bp
103,930,500 bp
103,930,520 bp
VCN3_pbsv.vcf
VCN3_HiFi_pbmm2.bam Coverage
[0 - 35]
VCN3_HiFi_pbmm2.bam
VCN3_Illumina_BWA.bam Coverage
[0 - 27]
VCN3_Illumina_BWA.bam
Sequence

### IS6_VCN4-VCN5_chr1:243273502

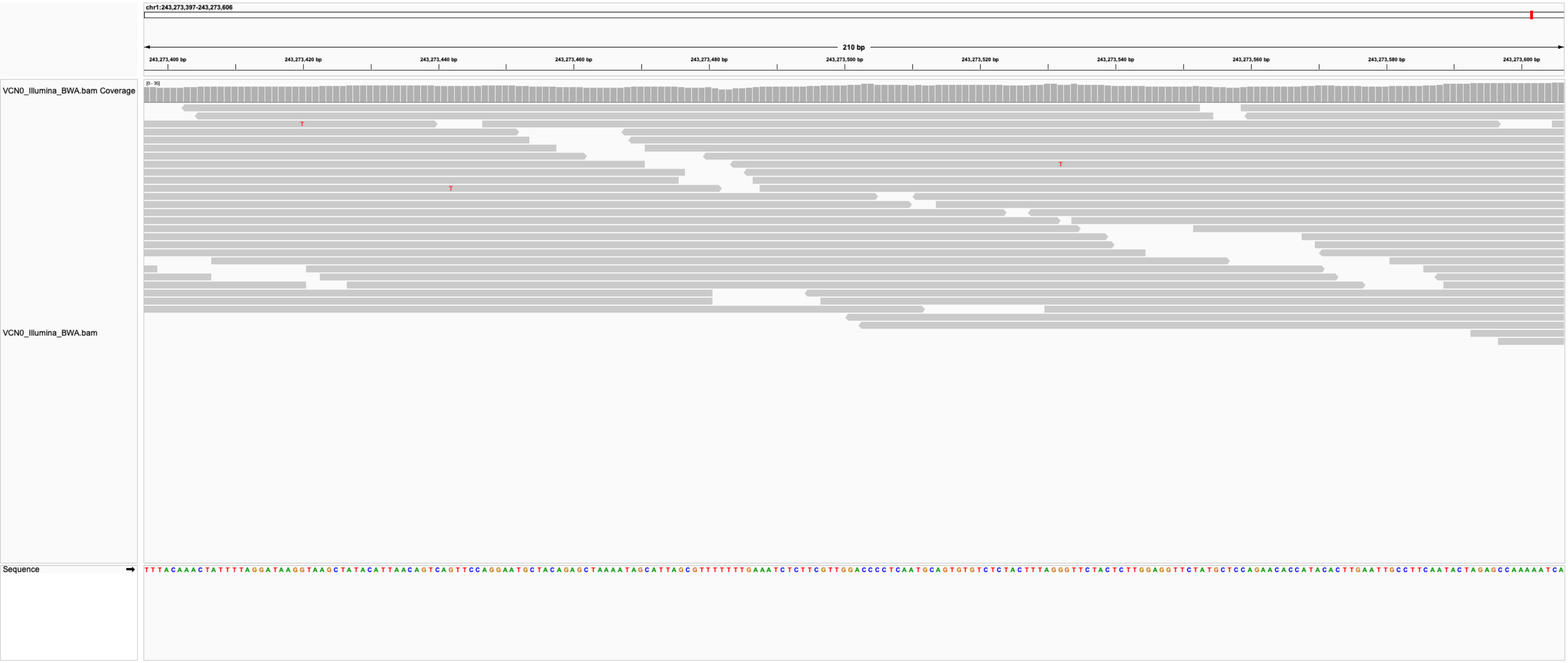

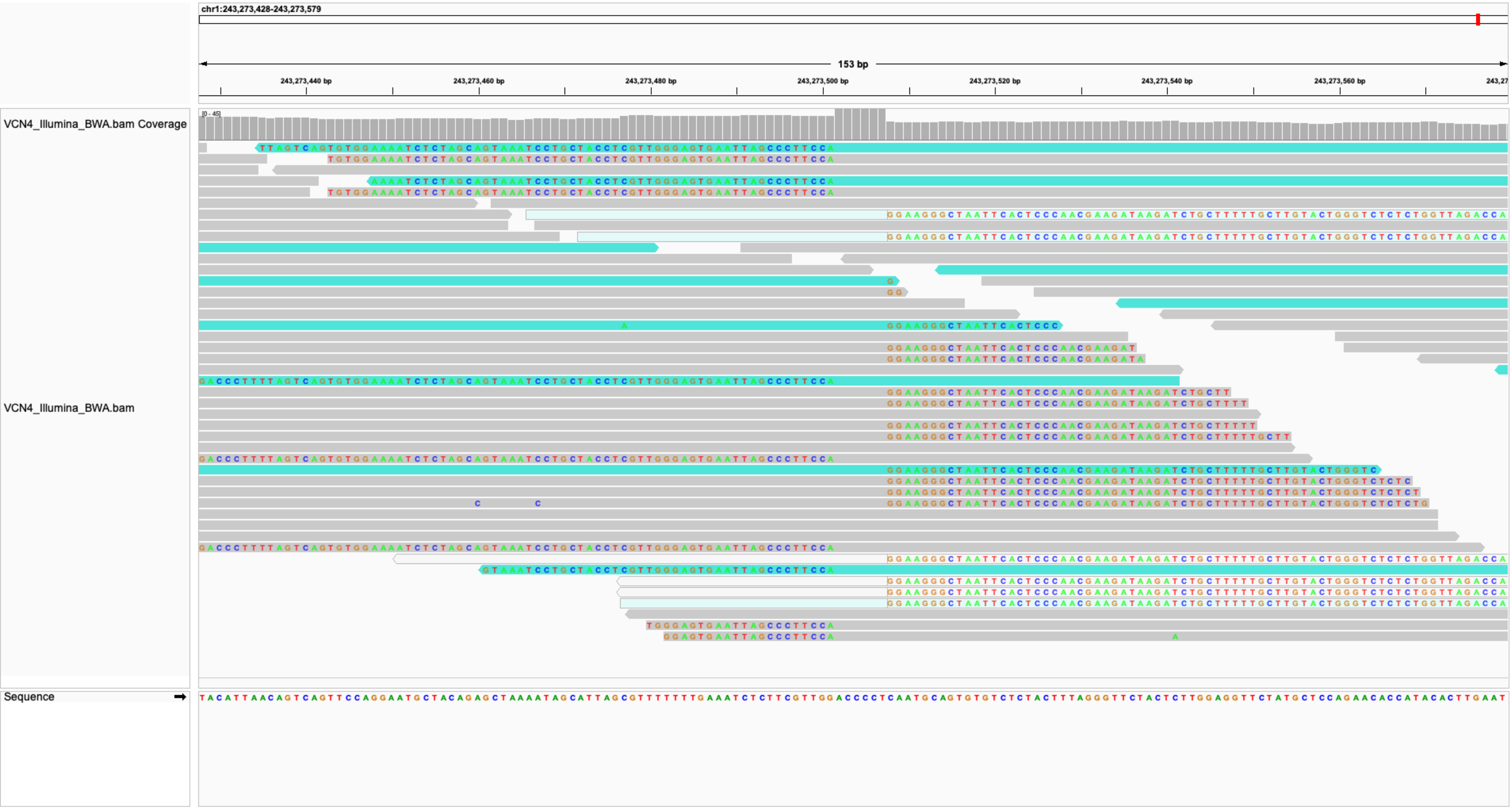
chr1:243,273,428-243,273,579
153 bp
243,273,440 bp
243,273,460 bp
243,273,480 bp
243,273,500 bp
243,273,520 bp
243,273,540 bp
243,273,560 bp
VCN4_Illumina_BWA.bam Coverage
[0 - 45]
VCN4_Illumina_BWA.bam
Sequence

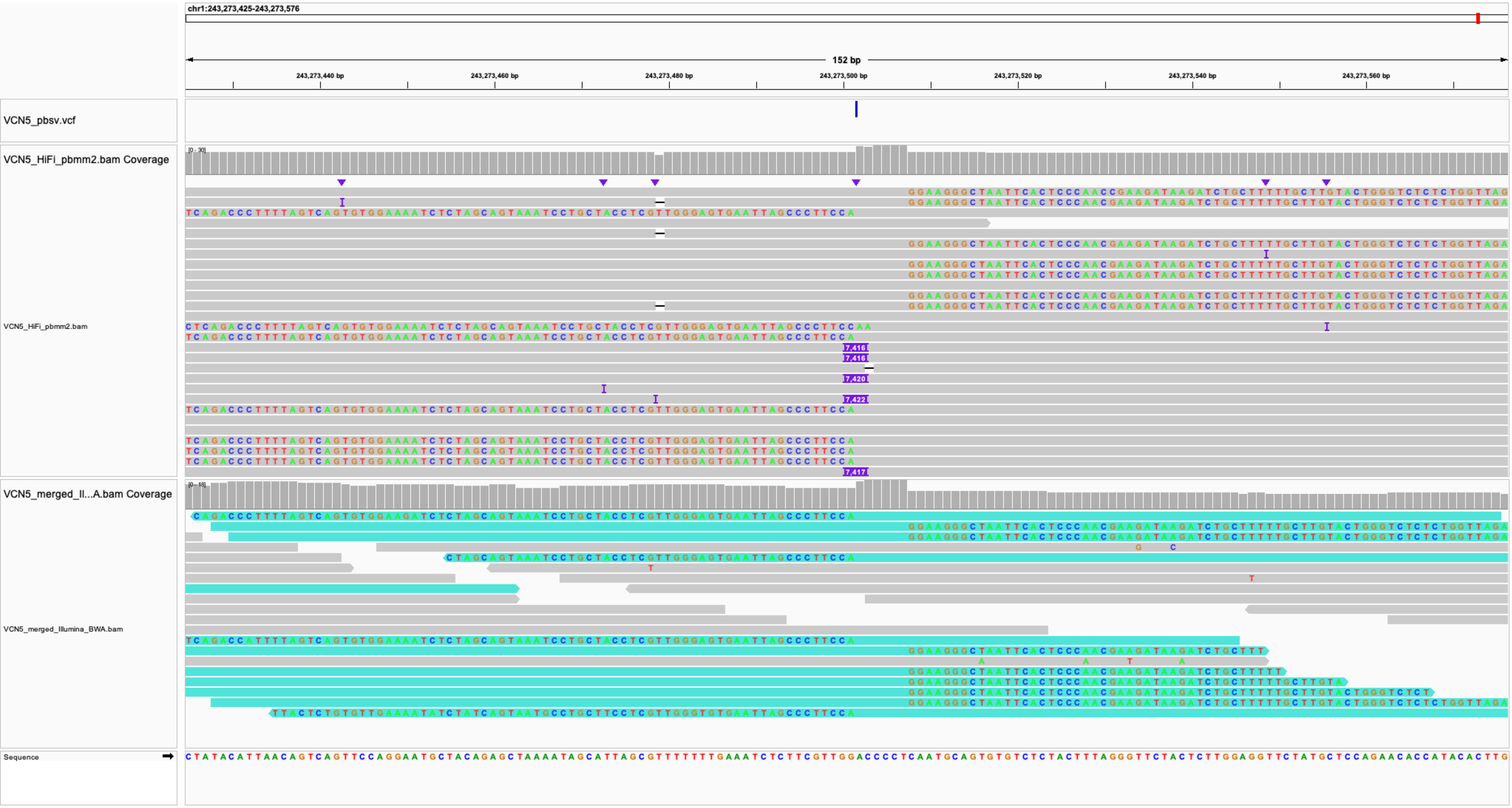
chr1:243,273,425-243,273,576
152 bp
243,273,440 bp
243,273,460 bp
243,273,480 bp
243,273,500 bp
243,273,520 bp
243,273,540 bp
243,273,560 bp
VCN5_pbsv.vcf
VCN5_HiFi_pbmm2.bam Coverage
[0 - 30]
VCN5_HiFi_pbmm2.bam
7,416
7,416
7,420
7,422
7,417
VCN5_merged_Il...A.bam Coverage
[0 - 18]
VCN5_merged_Illumina_BWA.bam
Sequence

### IS7_VCN4-VCN5_chr3:18115277

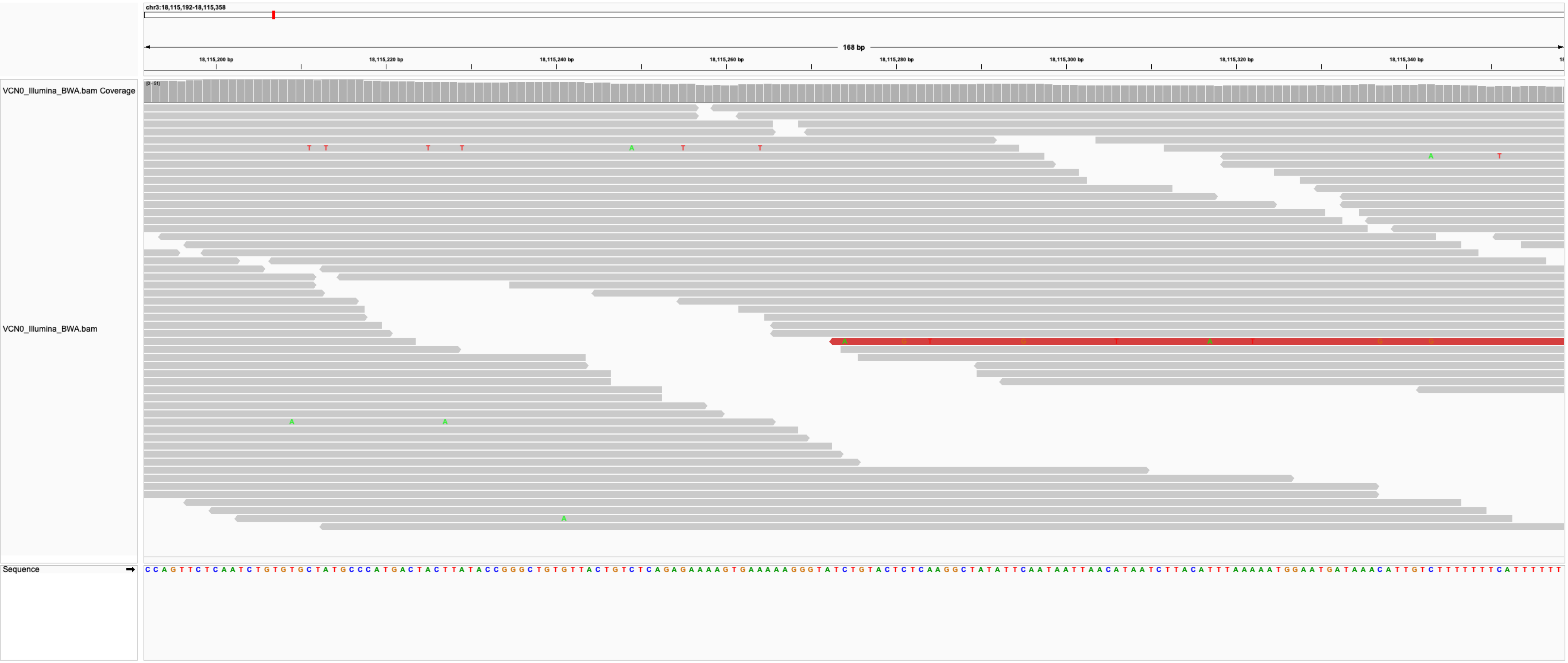

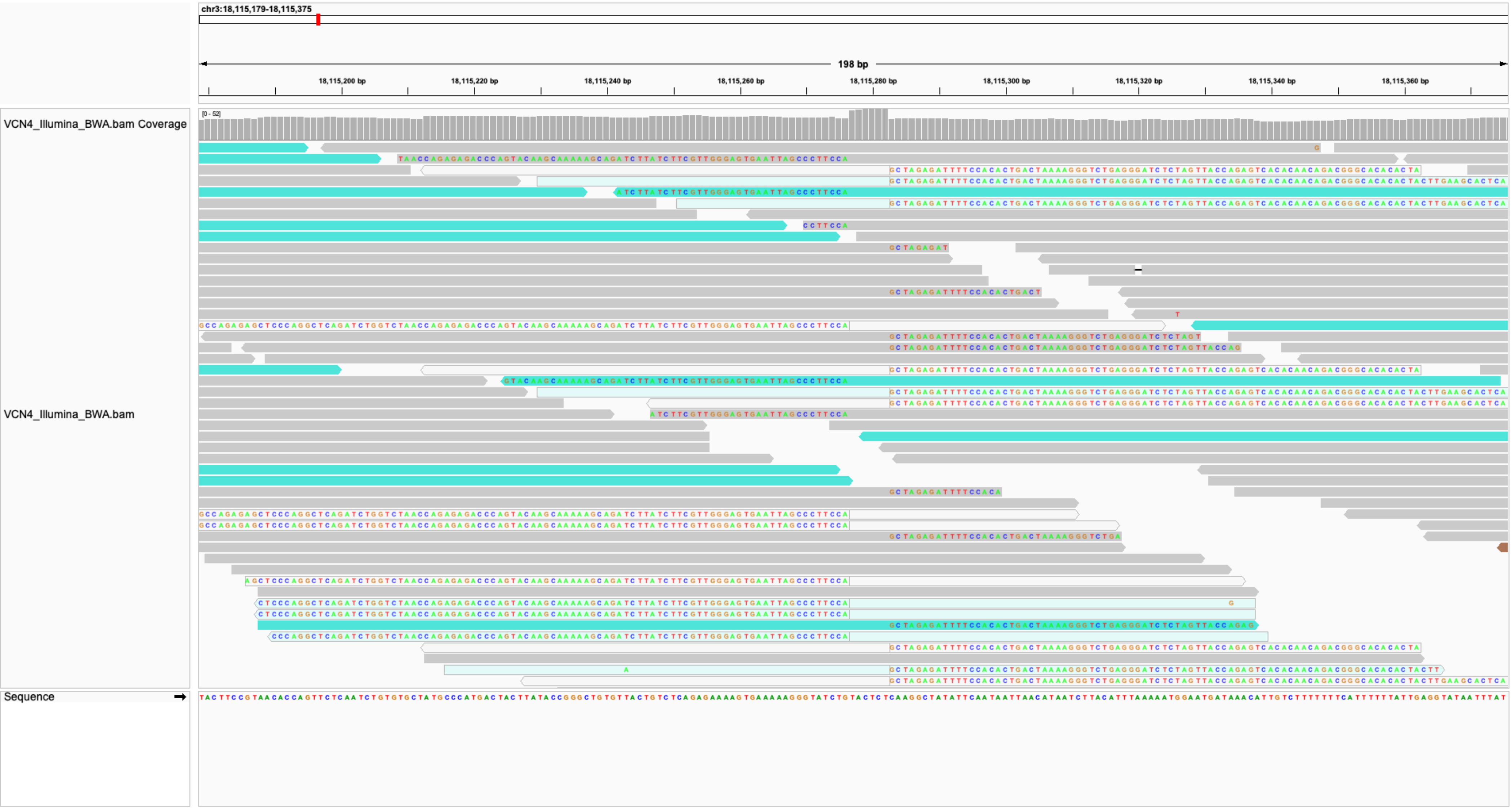

chr3:18,115,179-18,115,375
198 bp
18,115,200 bp
18,115,220 bp
18,115,240 bp
18,115,260 bp
18,115,280 bp
18,115,300 bp
18,115,320 bp
18,115,340 bp
18,115,360 bp
VCN4_Illumina_BWA.bam Coverage
[0 - 52]
VCN4_Illumina_BWA.bam
Sequence

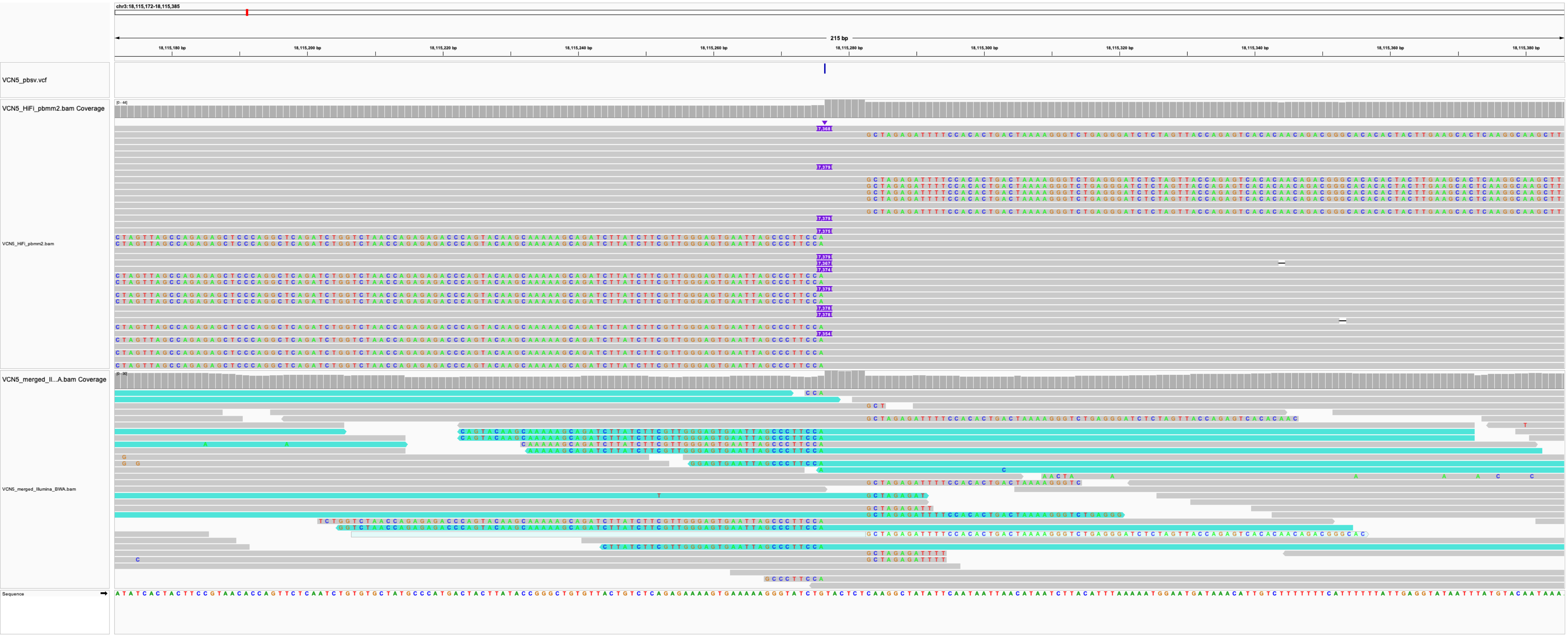

chr3:18,115,172-18,115,385
215 bp
VCN5_pbsv.vcf
VCN5_HiFi_pbmm2.bam Coverage
VCN5_HiFi_pbmm2.bam
VCN5_merged_Il...A.bam Coverage
VCN5_merged_Illumina_BWA.bam
Sequence

### IS8_VCN4-VCN5_chr6:15601040

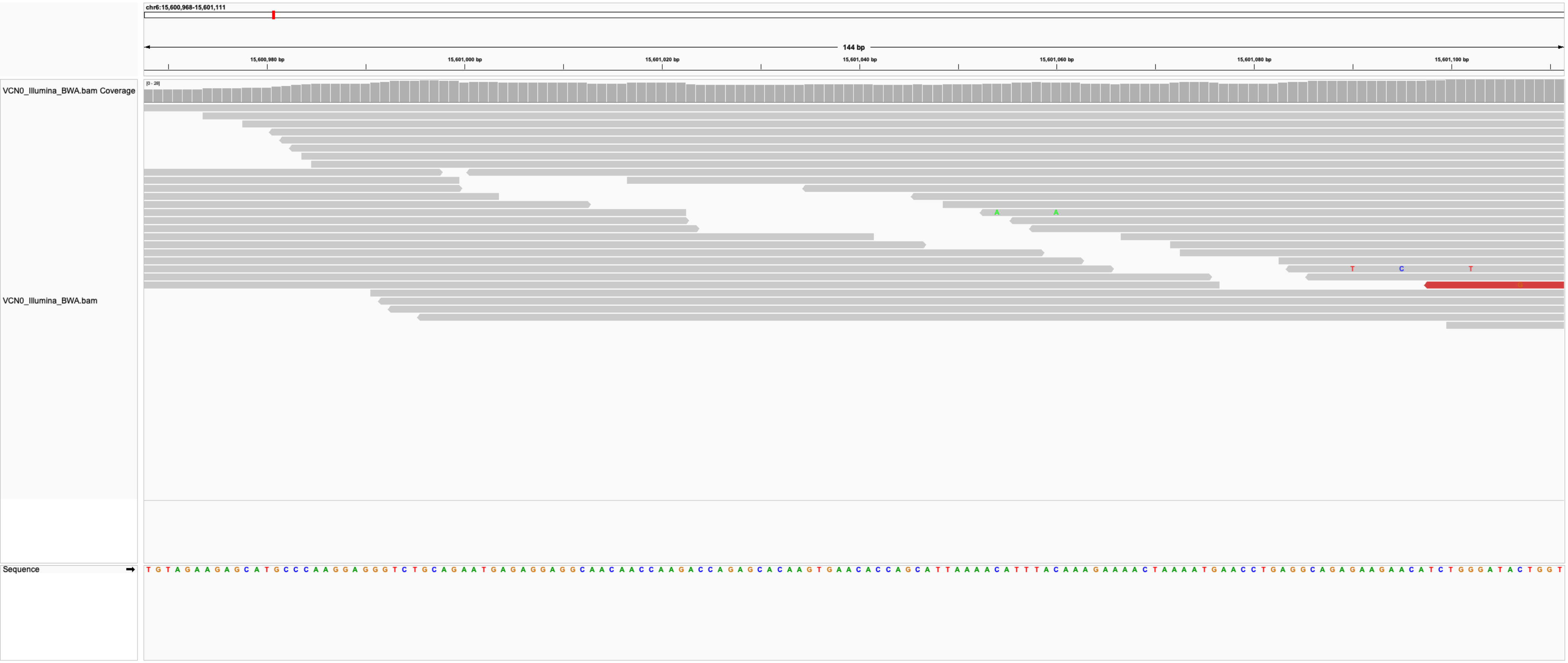

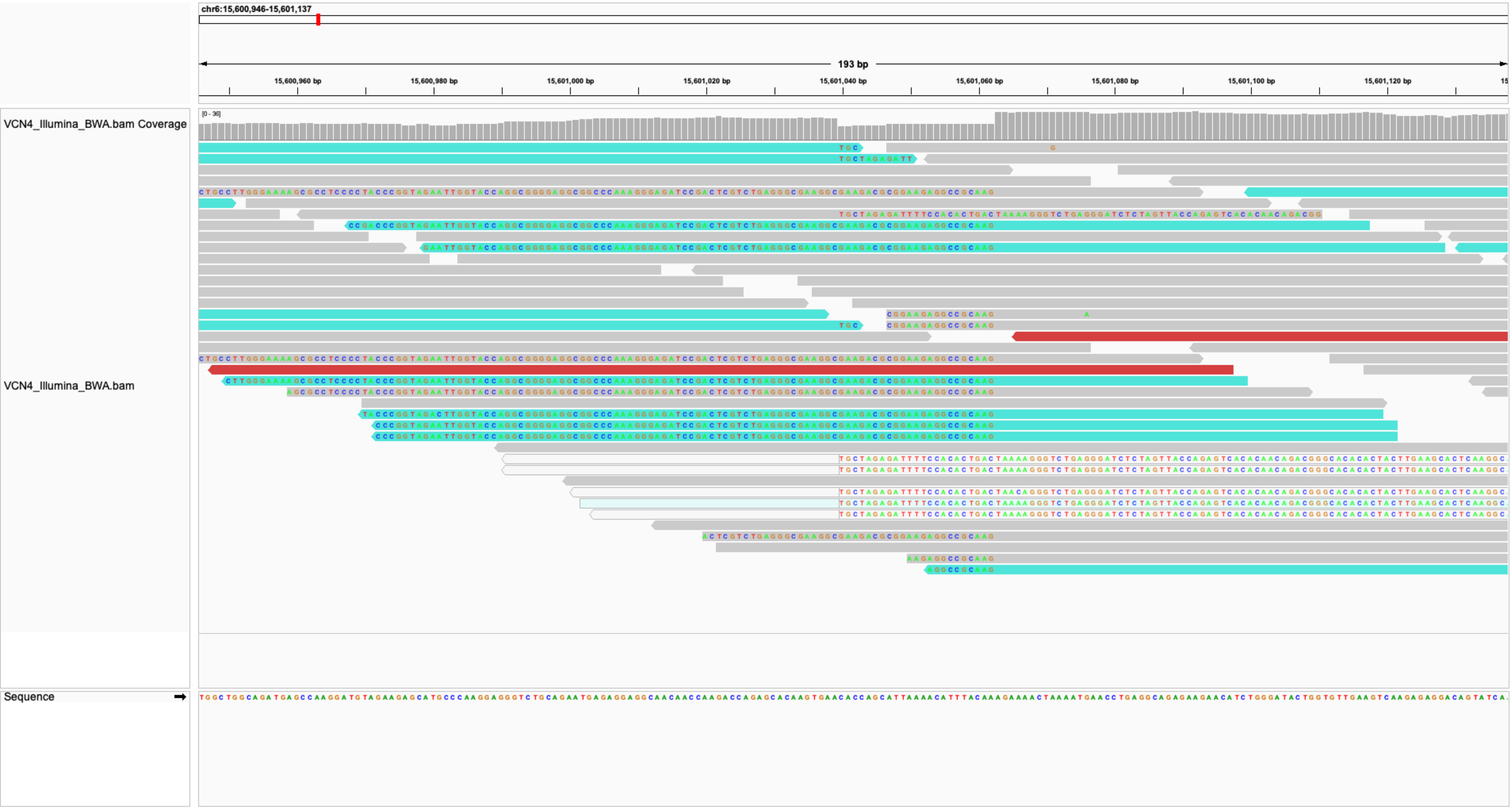
chr6:15,600,946-15,601,137
193 bp
15,600,960 bp
15,600,980 bp
15,601,000 bp
15,601,020 bp
15,601,040 bp
15,601,060 bp
15,601,080 bp
15,601,100 bp
15,601,120 bp
VCN4_Illumina_BWA.bam Coverage
[0 - 36]
VCN4_Illumina_BWA.bam
Sequence

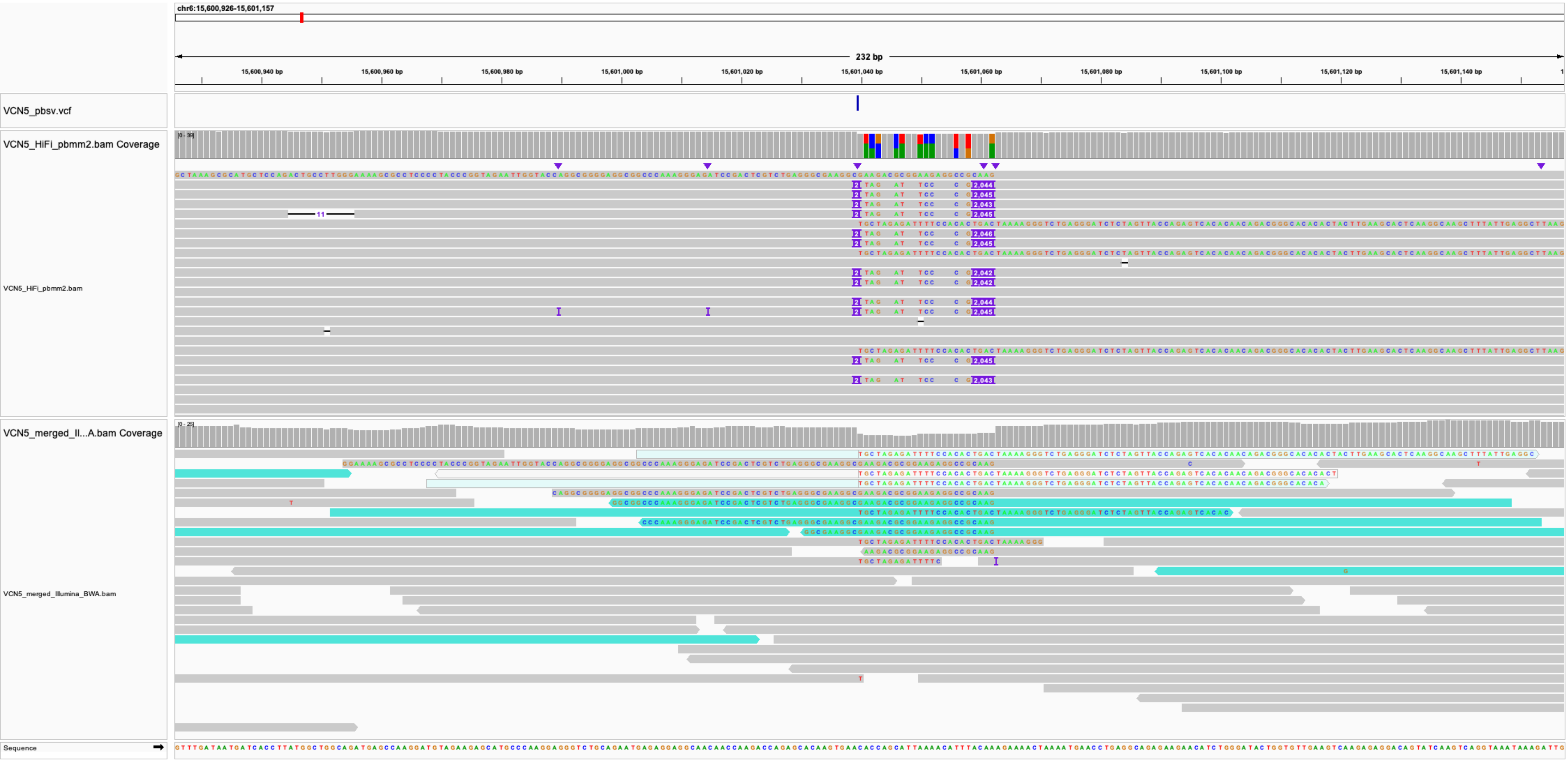
chr6:15,600,926-15,601,157
232 bp
15,600,940 bp
15,600,960 bp
15,600,980 bp
15,601,000 bp
15,601,020 bp
15,601,040 bp
15,601,060 bp
15,601,080 bp
15,601,100 bp
15,601,120 bp
15,601,140 bp
VCN5_pbsv.vcf
VCN5_HiFi_pbmm2.bam Coverage
VCN5_HiFi_pbmm2.bam
VCN5_merged_Il...A.bam Coverage
VCN5_merged_Illumina_BWA.bam
Sequence

### IS9_VCN4-VCN5_chr6:24825065

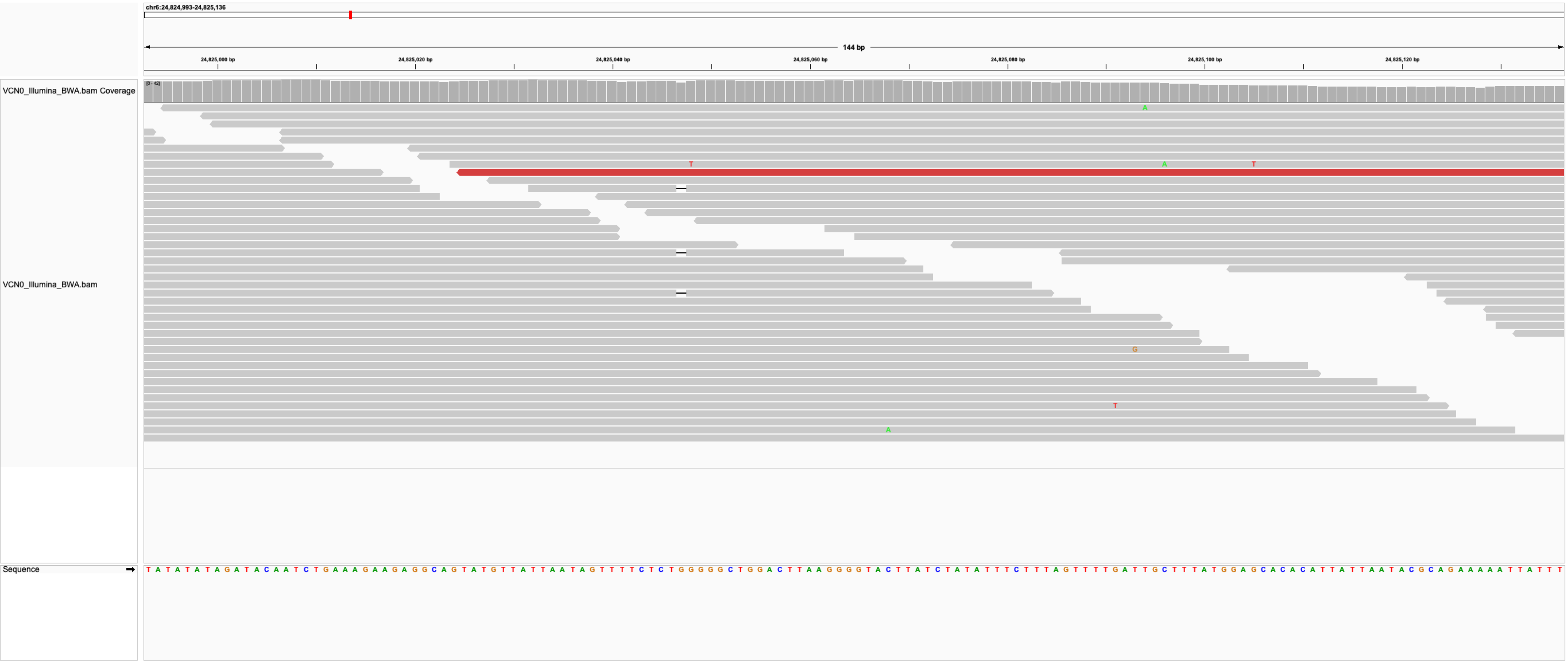

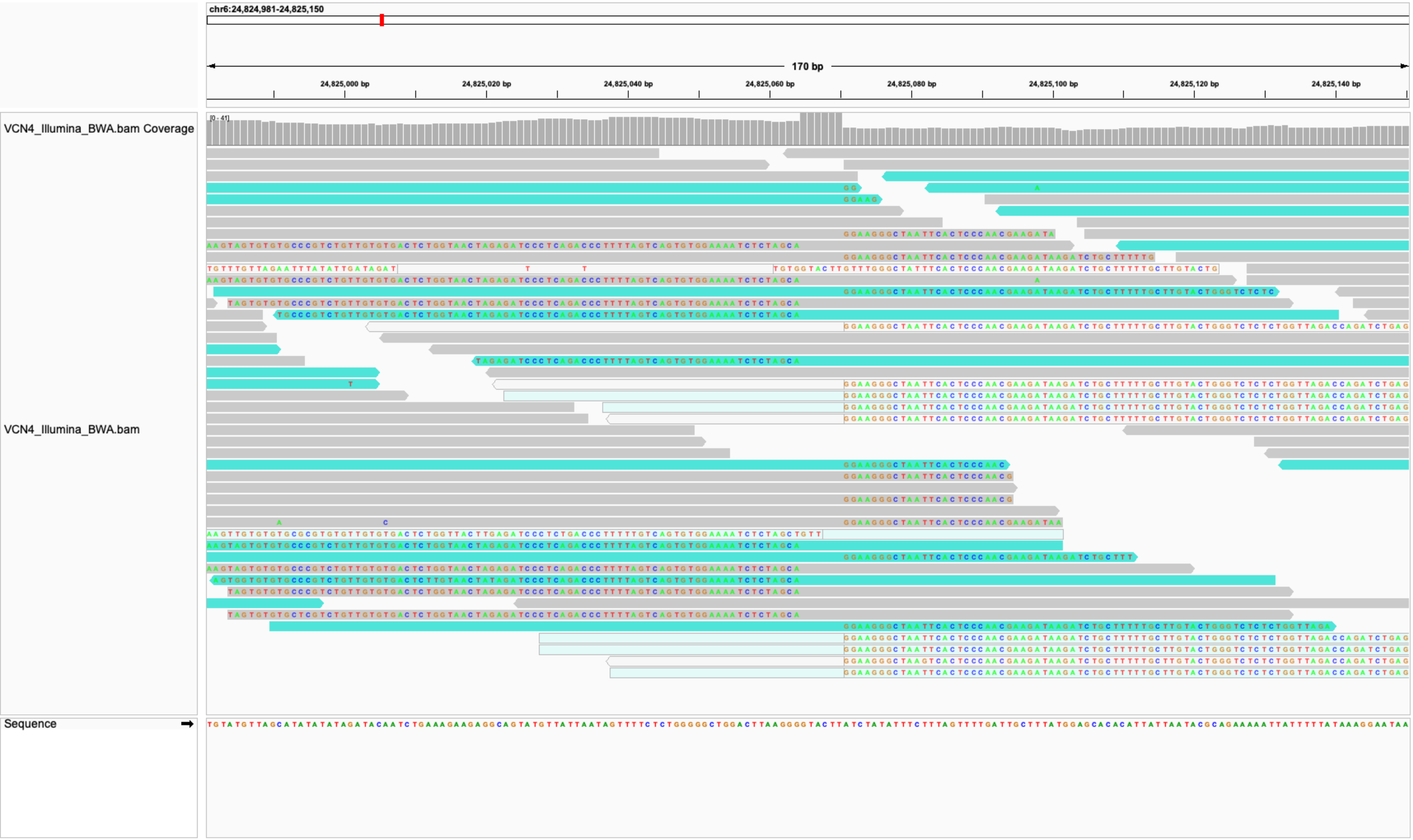
chr6:24,824,981-24,825,150
170 bp
24,825,000 bp
24,825,020 bp
24,825,040 bp
24,825,060 bp
24,825,080 bp
24,825,100 bp
24,825,120 bp
24,825,140 bp
VCN4_Illumina_BWA.bam Coverage
[0 - 41]
VCN4_Illumina_BWA.bam
Sequence

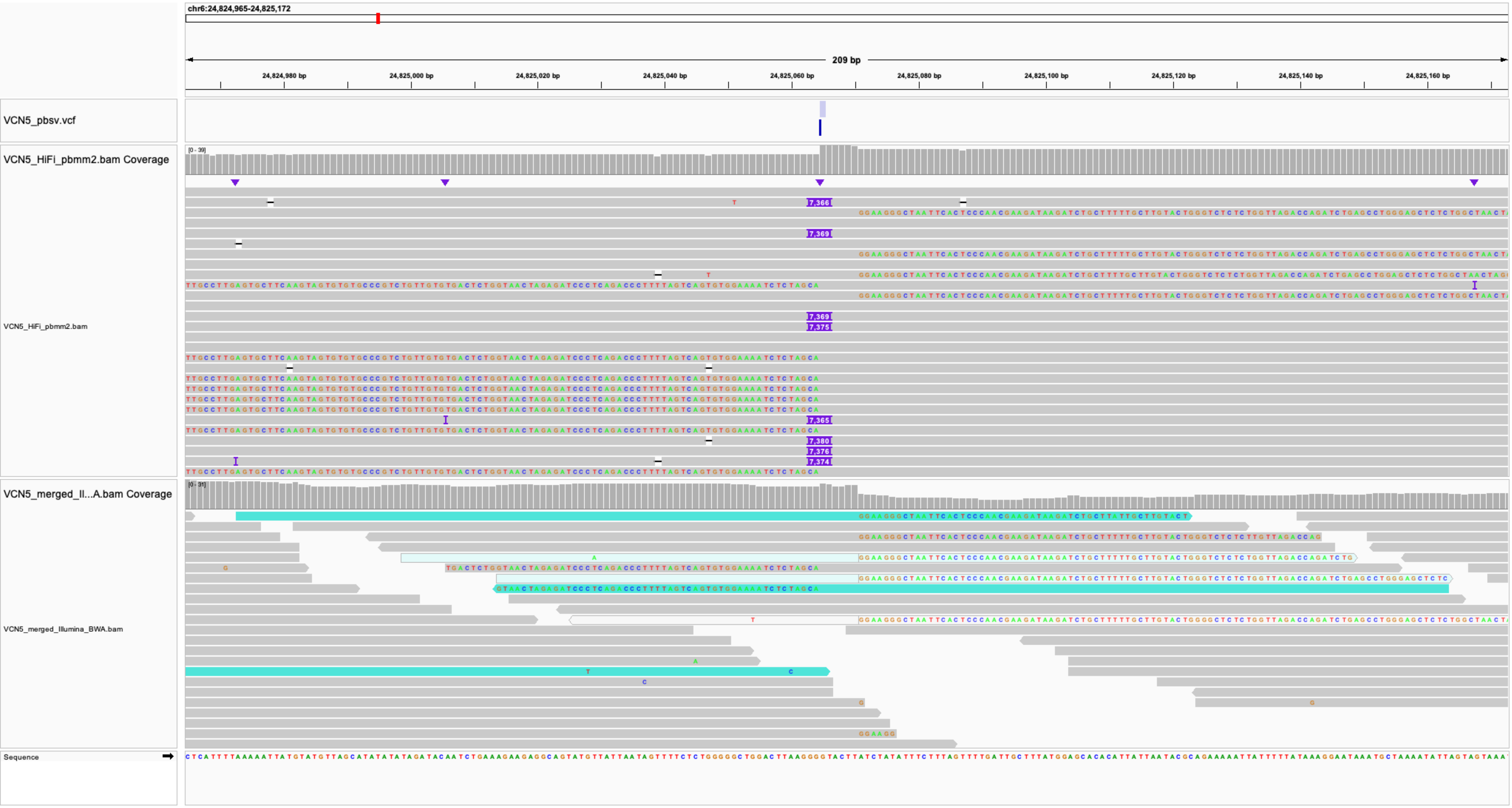
chr6:24,824,965-24,825,172
209 bp
24,824,980 bp
24,825,000 bp
24,825,020 bp
24,825,040 bp
24,825,060 bp
24,825,080 bp
24,825,100 bp
24,825,120 bp
24,825,140 bp
24,825,160 bp
VCN5_pbsv.vcf
VCN5_HiFi_pbmm2.bam Coverage
[0 - 39]
VCN5_HiFi_pbmm2.bam
7,366
7,369
7,369
7,375
7,365
7,380
7,376
7,374
VCN5_merged_Il...A.bam Coverage
[0 - 31]
VCN5_merged_Illumina_BWA.bam
Sequence

### IS10_VCN5_chr5:39044696

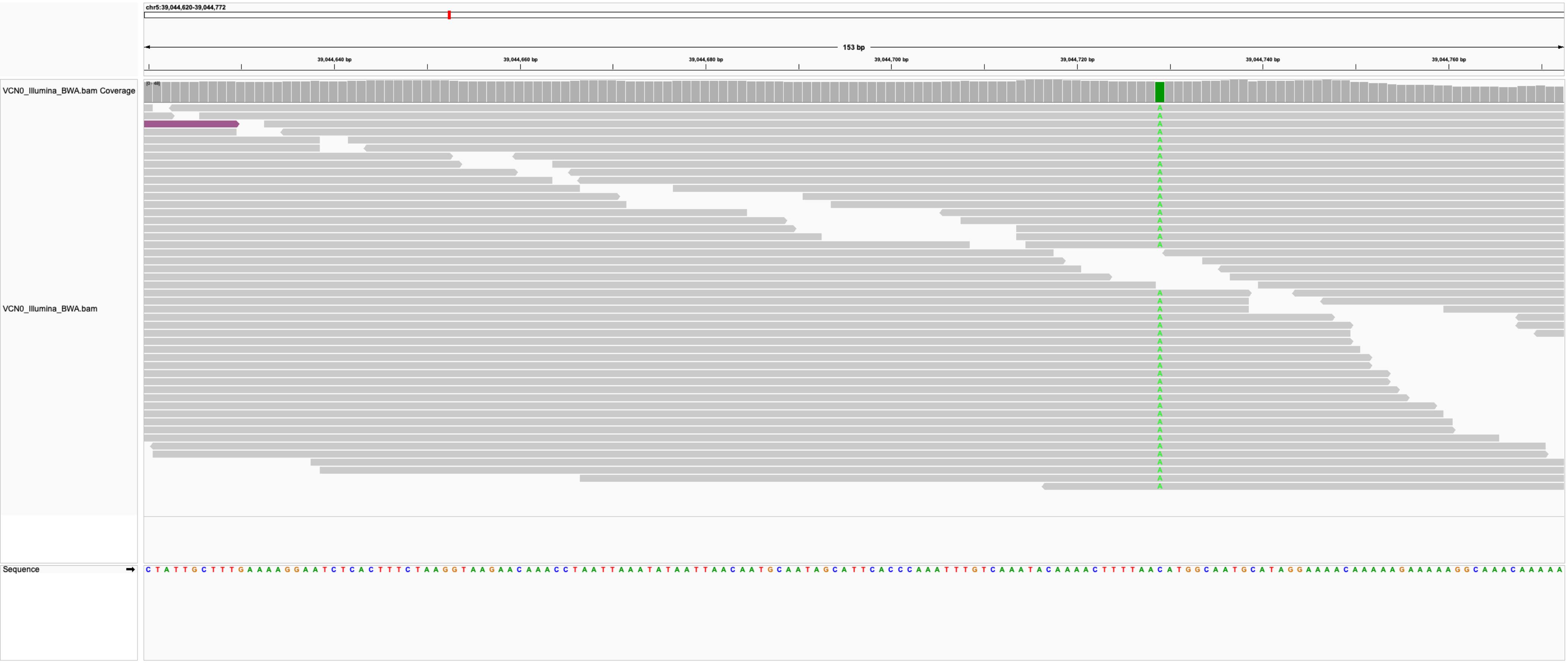

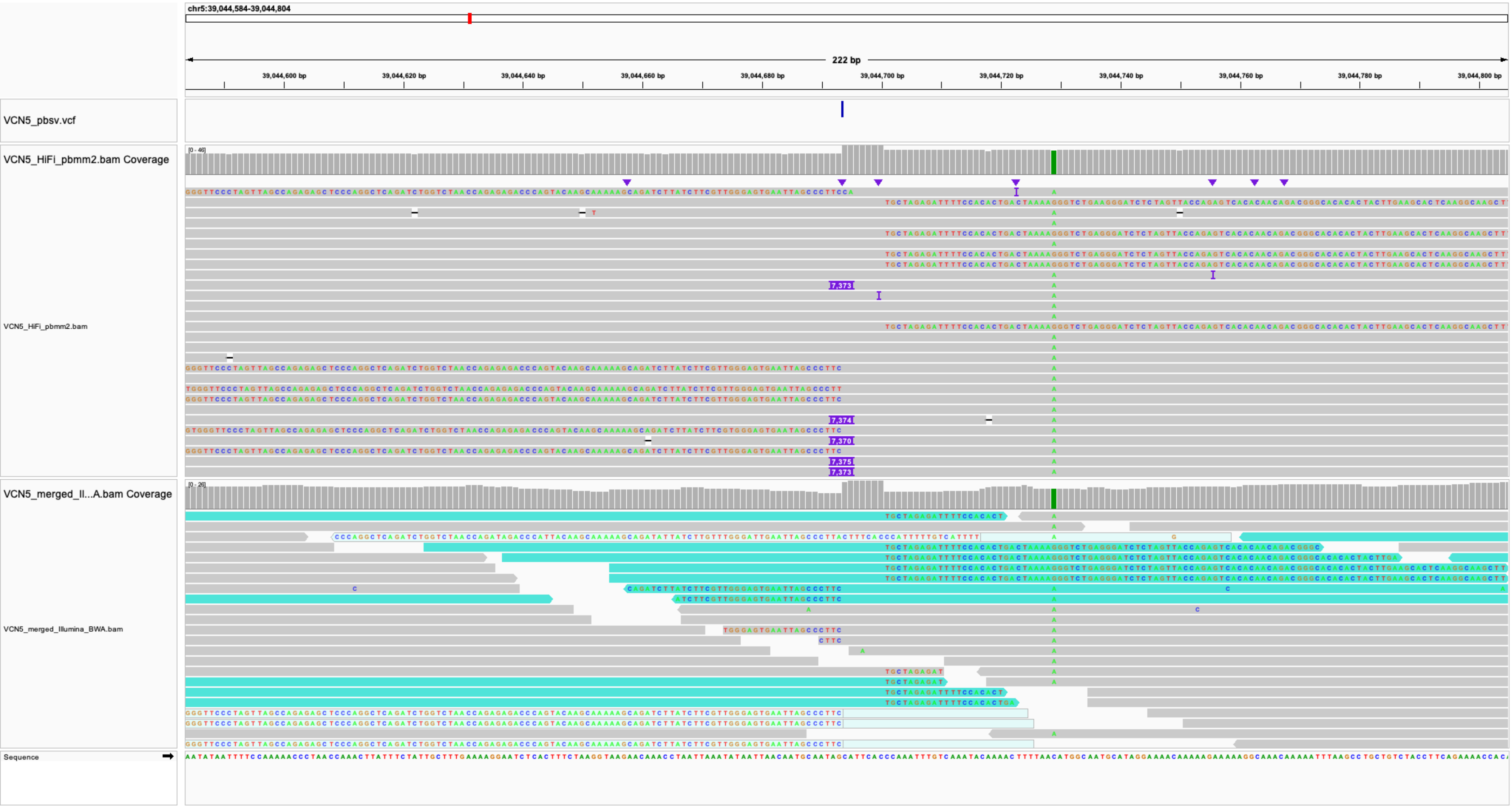
chr5:39,044,584-39,044,804
222 bp
39,044,600 bp
39,044,620 bp
39,044,640 bp
39,044,660 bp
39,044,680 bp
39,044,700 bp
39,044,720 bp
39,044,740 bp
39,044,760 bp
39,044,780 bp
39,044,800 bp
VCN5_pbsv.vcf
VCN5_HiFi_pbmm2.bam Coverage
[0 - 46]
VCN5_HiFi_pbmm2.bam
VCN5_merged_Il...A.bam Coverage
[0 - 26]
VCN5_merged_Illumina_BWA.bam
Sequence
